# Guidelines for Performing Ribosome Profiling in Plants Including Structural Analysis of rRNA Fragments

**DOI:** 10.1101/2023.11.16.567332

**Authors:** Michael K. Y. Ting, Yang Gao, Rouhollah Barahimipour, Rabea Ghandour, Jinghan Liu, Federico Martinez-Seidel, Julia Smirnova, Vincent Leon Gotsmann, Axel Fischer, Michael J. Haydon, Felix Willmund, Reimo Zoschke

## Abstract

Ribosome profiling (or Ribo-seq) is a technique that provides genome-wide information on the translational landscape (translatome). Across different plant studies, variable methodological setups have been described which raises questions about the general comparability of data that were generated from diverging methodologies. Furthermore, a common problem when performing Ribo-seq are abundant rRNA fragments that are wastefully incorporated into the libraries and dramatically reduce sequencing depth. To remove these rRNA contaminants, it is common to perform preliminary trials to identify these fragments because they are thought to vary depending on nuclease treatment, tissue source, and plant species. Here, we compile valuable insights gathered over years of generating Ribo-seq datasets from different species and experimental setups. We highlight which technical steps are important for maintaining cross experiment comparability and describe a highly efficient approach for rRNA removal. Furthermore, we provide evidence that many rRNA fragments are structurally preserved over diverse nuclease regimes, as well as across plant species. Using a recently published cryo-electron microscopy (cryo-EM) structure of the tobacco 80S ribosome, we show that the most abundant rRNA fragments are spatially derived from the solvent-exposed surface of the ribosome. The guidelines presented here shall aid newcomers in establishing ribosome profiling in new plant species and provide insights that will help in customizing the methodology for individual research goals.

## Introduction

Ribosome profiling was first described in 2009 (Ingolia et al., 2009) and since then has revolutionized our understanding of translation by providing genome-wide information about ribosome occupancy within translated regions. The method uses a ribonuclease treatment to degrade regions of mRNAs that are not protected by translating ribosomes. The remaining ribosome protected fragments (RPFs also called ribosome footprints) can be purified and examined by deep sequencing, which provides genome-wide information on the translational landscape. In addition, the position of the ribosome peptidyl site (P-site) can be computationally estimated for each RPF, thereby providing codon-level resolution of the translation activity. The original ribosome profiling technique has continually been improved, and it is often fine-tuned for individual species and tissues. Examples of recent and comprehensive descriptions of this technique are available for yeast and mice (Gerashchenko and Gladyshev, 2017), human and drosophila (Douka et al., 2022), and bacteria (Mohammad et al., 2019). In plants, translatomes have been assessed in several species including Arabidopsis (*Arabidopsis thaliana*) (Liu et al., 2013; Juntawong et al., 2014; Merchante et al., 2015; Hsu et al., 2016; Lukoszek et al., 2016; Chen et al., 2022), maize (Lei et al., 2015; Chotewutmontri et al., 2018), tomato (Wu et al., 2019b; Chiu et al., 2022), and rice (Yang et al., 2020; Yang et al., 2021). Since plant chloroplasts encode a distinct set of prokaryote-like ribosomes, specialized protocols also exist for assessing the chloroplast translatome (Zoschke et al., 2013; Scharff et al., 2017; Chotewutmontri and Barkan, 2018; Gawroński et al., 2018).

With the growing number of plant ribosome profiling studies, methodological variations have arisen, which might introduce potential biases at several steps (Bartholomäus et al., 2016). *Extraction buffer composition*: the ionic strength and buffering capacity of the extraction buffer can affect the observed behavior of RPFs (Hsu et al., 2016). *Choice of ribonuclease*: several ribonucleases have been used for the generation of RPFs, including RNase I, A, T1 and MNase (Gerashchenko and Gladyshev, 2017). Some ribonucleases exhibit preferential cleavage at specific motifs, thereby confounding codon resolution. To date, the most widely used ribonuclease for Ribo-seq in eukaryotes is RNase I (Ingolia, 2016), whereas MNase is the preferred ribonuclease for Ribo-seq in prokaryotes (Mohammad et al., 2019). *Ribonuclease treatment*: the amount of ribonuclease, digestion time, and digestion temperature can vary across studies. In addition, ribonuclease treatment can be performed directly on cell lysates or on purified polysomes. *RPF purification strategy*: some protocols capture RPFs within a narrow size range (e.g., 28-30 nt), which enriches the highly periodic RPFs (Hsu et al., 2016). Others prefer to use a broader size range (e.g., 20-40 nt), which also has notable benefits (Chotewutmontri et al., 2018). Importantly, a broader size range is inclusive of unique RPFs that convey valuable information about translational dynamics, such as the 21 nt RPFs that represent ribosomes lacking a tRNA in the A-site (Wu *et al*., 2019a). *rRNA removal*: since ribosomes are composed of RNA, ribonuclease treatment unavoidably leads to the generation of widespread nicks in rRNA, creating fragments which can co-purify with RPFs. These unwanted rRNA fragments are wastefully incorporated into the sequencing libraries and substantially reduce the number of informative reads. Small scale sequencing tests are often performed to identify the major fragments from individual experimental setups (Mahboubi et al., 2021). Enzymatic strategies to remove rRNA have been described (Chung et al., 2015) but these methods have been shown to perturb codon-resolution (Zinshteyn et al., 2020). The most commonly applied approach to remove rRNA contamination is subtractive hybridization using biotinylated DNA oligonucleotides (oligos). *Library preparation*: the original ribosome profiling method used RNA circularization to incorporate RPFs into a cDNA library, which is a method still used by many labs. Libraries can also be prepared from kits designed for sequencing of small-RNA that utilize RNA ligases for adapter incorporation (Chotewutmontri et al., 2018), as well as ligation-free approaches that utilize polyadenylation and reverse transcription template-switching (Hornstein et al., 2016).

Such methodological variation can seem overwhelming to those performing ribosome profiling for the first time, and/or to those who wish to establish the technique in a new plant species. It also raises concerns of the comparability of datasets across different studies that have utilized different methodologies. Here, we focus on data reproducibility by compiling valuable insights gathered over years of generating Ribo-seq datasets from different plant species and experimental setups. We also provide a structural analysis of the rRNA fragments that regularly contaminate Ribo-seq libraries, and reveal patterns that are spatially preserved over diverse nuclease treatments, as well as across plant species. Overall, these guidelines are anticipated to be a valuable resource for the plant community and should be applicable to any Ribo-seq methodology.

## Materials and Methods

The following section provides information for the samples, which were prepared over different stages of protocol optimization. Thus, the data presented are derived from different plant material from diverse experiments. The detailed, fully optimized protocol is provided in the Supplemental Methods.

### Plant material

The tissue used for the comparison of RNase I and MNase digestion, was derived from 8-day old Arabidopsis seedlings (Col-0) grown on ½ Murashige and Skoog medium (Murashige and Skoog, 1962) with 6.8% agar and 1% sucrose, grown at 100 µmol m^-2^s^-1^ for 16h/8h light/dark cycles at 20 °C. The tissue used for refining RNase I treatment, rRNA depletion and comparison of ligation-free to ligation-based strategies, were derived from 14-day old Arabidopsis seedlings (Col-0), grown on ½ Murashige and Skoog media with 1% agar, grown at 100 µmol m^-2^s^-1^ for 12h/12h light/dark cycles at 20 °C. Tobacco (*Nicotiana tabacum*) tissue used for ribosome profiling was derived from a temperature-shift experiment, from leaves harvested from 28-day old plants grown on soil at 350 µmol m^-2^s^-1^ in 16h/8h light/dark cycles at 12 °C. Tobacco (*Nicotiana tabacum*) tissue used for polysome profiling (Figure 1) was derived from leaves harvested from 21-day old plants grown on soil at 350 µmol m^-2^s^-1^ in 16h/8h light/dark cycles at 24 °C.

**Figure 1.**
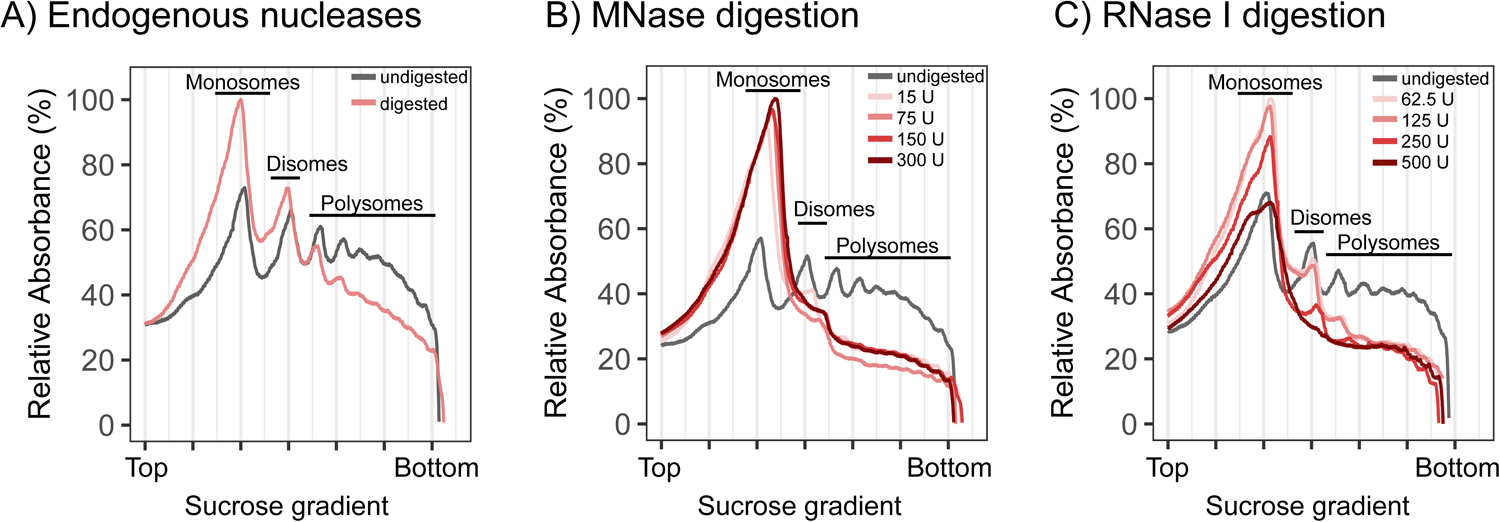
Qualitative assessment of ribonuclease concentrations by polysome profiling. Polysome profiling was used to determine optimal nuclease concentrations for converting polysomes into monosomes. **(A-C)** Sucrose gradient profiles of ribonucleoprotein particles derived from wild-type tobacco leaf lysates, following treatment with (A) Endogenous nucleases, or endogenous nucleases supplemented with (B) MNase or (C) RNase I. Undigested samples contain the ribonuclease inhibitor heparin to inhibit nuclease activity. Ribonucleoprotein particles were size-separated in sucrose density gradients (15, 30, 45, and 60% w/v from top to bottom) by ultracentrifugation as previously described (Gao et al., 2022). The signals of monosomes, disomes and polysomes were detected by UV absorbance measurements (254 nm). U, Units of applied ribonuclease.

### RNA and RPF isolation

Total RNA and RPFs were isolated as previously described (Trösch et al., 2018) with modifications described in the Supplemental Methods. The units (U) of ribonuclease used in this study are normalized to one mL of plant lysate, derived from 100 mg of plant fresh weight. Since Ca^2+^ is a known cofactor of MNase, samples digested with MNase include 5 mM CaCl_2_. All RPFs that were not rRNA-depleted were size-selected between 20-50 nt. All rRNA-depleted RPFs were size-selected between 20-35 nt. Details of rRNA depletion are available in the supplemental methods.

### Library preparation

For the ligation-free strategy, rRNA depleted RPFs were directly used as input for the D-plex small RNA-seq kit (Diagenode cat#C05030001), according to manufacturer’s instructions. Diagenode libraries are typically amplified with 7-9 PCR cycles. For the RNA-ligase strategy, the terminal ends of the RPFs were first repaired using T4 polynucleotide kinase (PNK; ThermoFisher, cat#EK0031). This was carried out in 20 µL volume with ∼100 ng of RPFs (un-depleted) or ∼30 nt of RPFs (rRNA-depleted), as described in the supplemental methods. After treatment, RPFs were directly used as input into the NEXTflex small RNA-seq kit v3 (Perkin Elmer, cat# NOVA-5132-06) or V4 (Perkin Elmer, cat#NOVA-5132-31), according to the manufacturer’s instructions. NEXTflex libraries are typically amplified using 14-16 PCR cycles. Libraries were sequenced on a Nextseq500 (SE75) or Novaseq6000 (SE100). The sequencing data have been deposited in NCBI’s Gene Expression Omnibus under accession number GSE226508.

### Identification of major rRNA fragments

To identify the most abundant rRNA fragments, pioneer Ribo-seq libraries were aligned to rRNA genes as described in the Supplemental methods. Each rRNA gene was then visually inspected in the IGV browser (http://software.broadinstitute.org/software/igv) to identify regions with high coverage that were repeatedly present in the majority of the libraries. Complementary biotinylated DNA oligos were designed (Table S1 and S2) and mixed together in molar ratios equivalent to the relative averaged abundance of the target rRNA contaminant within these pioneer libraries.

### Mapping rRNA fragments to the ribosome structure

The reference structure used in this work corresponds to the translating cytosolic ribosome of *Nicotiana tabacum* (PDB: 8B2L, EMDB: 15806) (Smirnova et al., 2023). The structure was solved by using single-particle cryo-electron microscopy to an overall resolution of 2.2 Å. The molecular model of the tobacco 80S ribosome contains in total 91% of the rRNA residues within the small and 95% of the rRNA residues within the large ribosomal subunit. The top contaminating rRNA fragments, derived from pioneer tobacco Ribo-seq datasets, were mapped to the 80S ribosome model using PyMOL (The PyMOL Molecular Graphics System, Version 1.2r3pre, Schrödinger, LLC) and colored according to their relative abundance.

## Results and Discussion

### Effects of variable nuclease treatments

In the early stages of establishing plant ribosome profiling in our group, we were initially concerned that RPFs generated using different nucleases and/or nuclease concentrations, might lead to technical variation that limits reproducibility. For example, not using sufficient nuclease (under digestion) could become problematic if there exists a population of ribosomes that preferentially remained in the polysome fraction (i.e., some transcripts that are more resistant to nuclease digestion due to RNA binding proteins or RNA secondary structure). Such a bias would result in reduced RPF yield, and/or alter the quantitative translatome. On the other hand, using excessive nuclease (over digestion) was anticipated to increase rRNA fragmentation, overly breaking down ribosomes and subsequently reducing RPF yield. To address these concerns, polysome profiling was performed to identify the minimal nuclease concentration required to efficiently convert polysomes into monosomes, without causing excessive monosome breakdown. When performing digestion directly on cell lysate, endogenous nuclease activity also contributes to monosome formation (Figure 1A). All MNase concentrations tested produced similar profiles, highlighting the general robustness of treatments with this nuclease (Figure 1B). For RNase I, disomes and higher order polysomes remained visible from samples treated with less than 250 U, suggestive of under digestion, whereas using more than 250 U resulted in monosome reduction, suggestive of over digestion (Figure 1C).

To expand on the polysome profile observations, 6 of the nuclease treatments were selected for Ribo-seq. As expected, rRNA dominates the library composition (Figure 2A). Importantly, a crosswise comparison of RPF density over annotated genes displayed high correlations across all datasets (Figure 2B), indicating that translatome data generated using MNase and RNase I are quantitatively comparable over a wide range of digestion conditions. These observations support the fair comparison of published datasets generated using different nuclease regimes, which is particularly relevant when attempting to integrate plant translatome data from chloroplast-focused studies that use MNase, to nuclear-focused studies that use RNase I.

**Figure 2.**
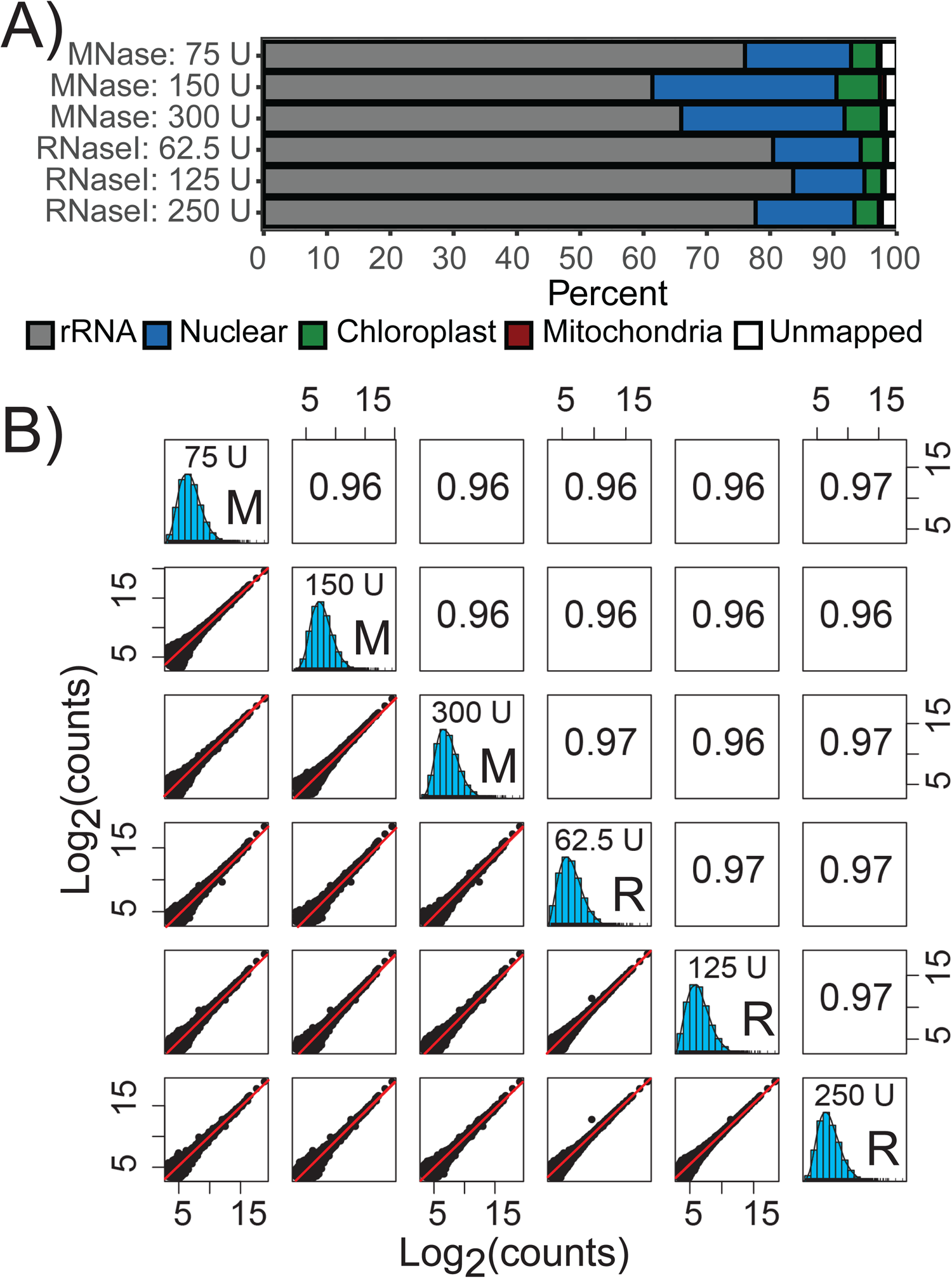
Quantitative comparison of Ribo-seq datasets generated over diverse nuclease treatments. **(A)** Ribo-seq library composition, based on RPF mapping location. **(B)** Spearman’s correlation of RPF density over annotated protein-coding sequences (CDS). Lowly translated genes with fewer than 10 counts were excluded from the analysis. The line of best fit is shown as the red diagonal. M, MNase-treated; R, RNase I-treated; U, Units of applied ribonuclease.

Next, the qualitative properties of each translatome were assessed. True RPFs should predominantly be found in annotated protein-coding sequences (CDS), which was indeed the case for all treatments (Figure 3A). This was also confirmed through manual inspection of RPF density over selected genes (Figure S1). Since RPF density is reflective of translational kinetics, increased RPF density should be visible every 3 nucleotides as ribosomes slow down to decode each codon (Ingolia, 2016). This pattern is commonly referred to as triplet periodicity and is often utilized as a quality measure and for the statistical detection of actively translated reading frames (Calviello et al., 2015; Raj et al., 2016; Xiao et al., 2018; Xu et al., 2018; Choudhary et al., 2020). To measure triplet periodicity, RPFs were positioned at their P-site and the RPF density was quantified over the three frames of translation. Periodicity was only observed for samples treated with RNase I (Figure 3B,C), reaffirming the previously described qualitative benefits of using RNase I over MNase (Gerashchenko and Gladyshev, 2017).

**Figure 3.**
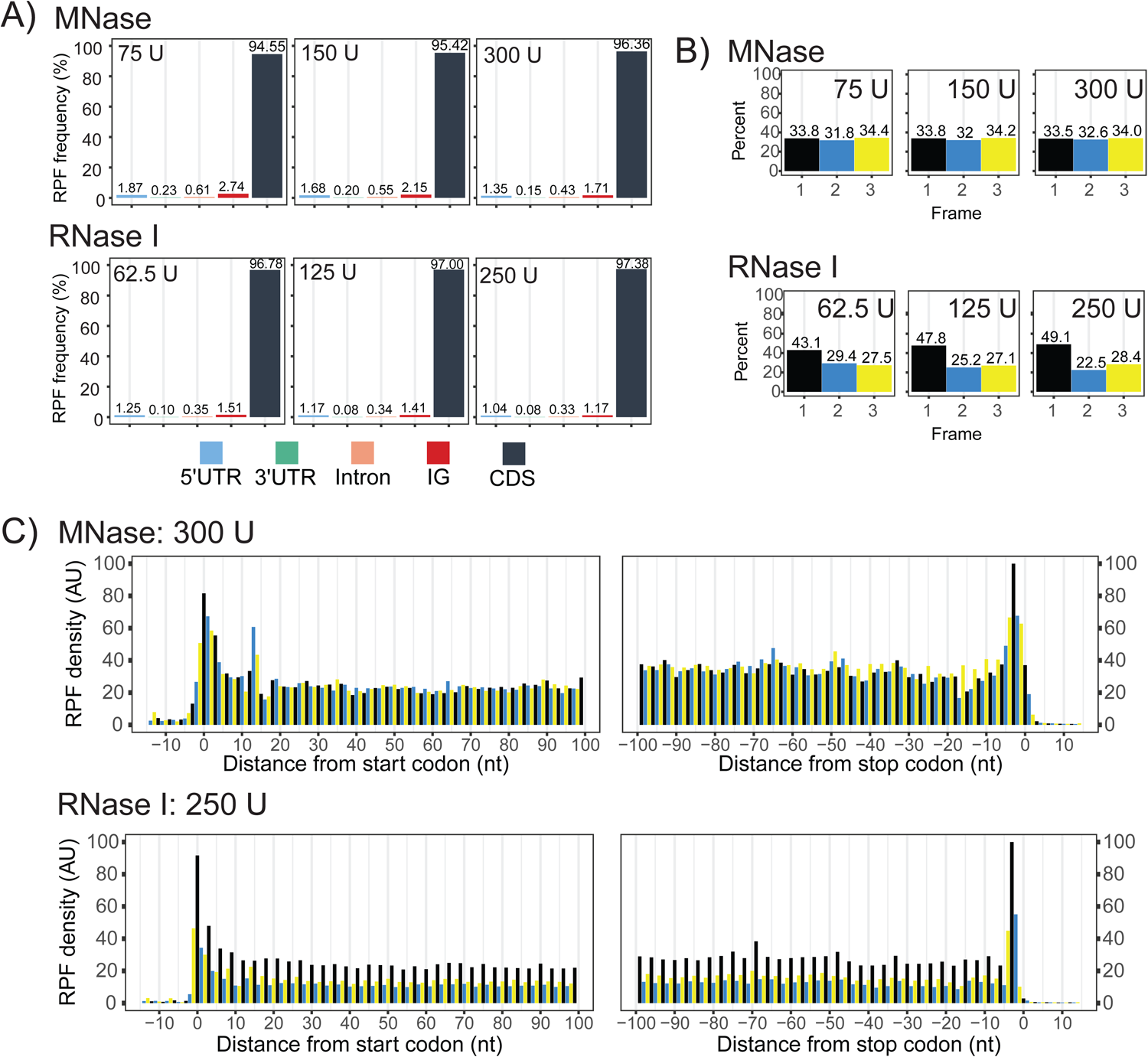
Qualitative comparison of Ribo-seq datasets generated over diverse nuclease treatments. **(A)** Proportion of mapped RPFs across genomic features. The percentages are calculated after removal of rRNA. **(B)** Quantification of RPFs in each frame of translation. Frame 1 is in reference to the annotated start and stop codons. **(C)** Metagene analysis for samples generated following digestion with 300 U of MNase, or 250 U of RNase I. RPFs are positioned at their P-site using offsets calculated from the 5’-position. U, Units of applied ribonuclease; UTR, untranslated regions; IG, intergenic regions; CDS, annotated protein-coding sequences.

In plants, most cytosolic RPFs are reported to be ∼28-29 nt in size which are characterized by strong triplet periodicity (Hsu et al., 2016; Chotewutmontri et al., 2018). RPFs larger than 28-29 nt tend to display lower triplet periodicity, which may be attributed to excess nucleotide(s) at either the 5’ or 3’ end (Figure 4A) thereby confounding P-site estimations. Under the conditions tested, the majority of cytosolic RPFs in our samples were larger than 29 nt, indicating under digestion. However, the expected shift towards smaller sizes was observed as nuclease concentration was increased (Figure 4B,C). This size shift was not observed for the rRNA, indicating robust protection of specific rRNA fragments. In addition, a secondary cytosolic RPF peak at ∼20-24 nt was present in the samples, being more prominent with RNase I treatment (Figure 4C). This secondary peak likely corresponds to ribosomes with an empty A-site (Wu *et al*., 2019a), illustrating that diverse species of RPFs are captured, and highlighting the importance of using a broader size selection. Two peaks were also observed for chloroplast RPFs, which is a pattern also described in maize (Chotewutmontri and Barkan, 2016). Additional analysis of these two populations are provided below in a specific section concerning chloroplast-derived RPFs.

**Figure 4.**
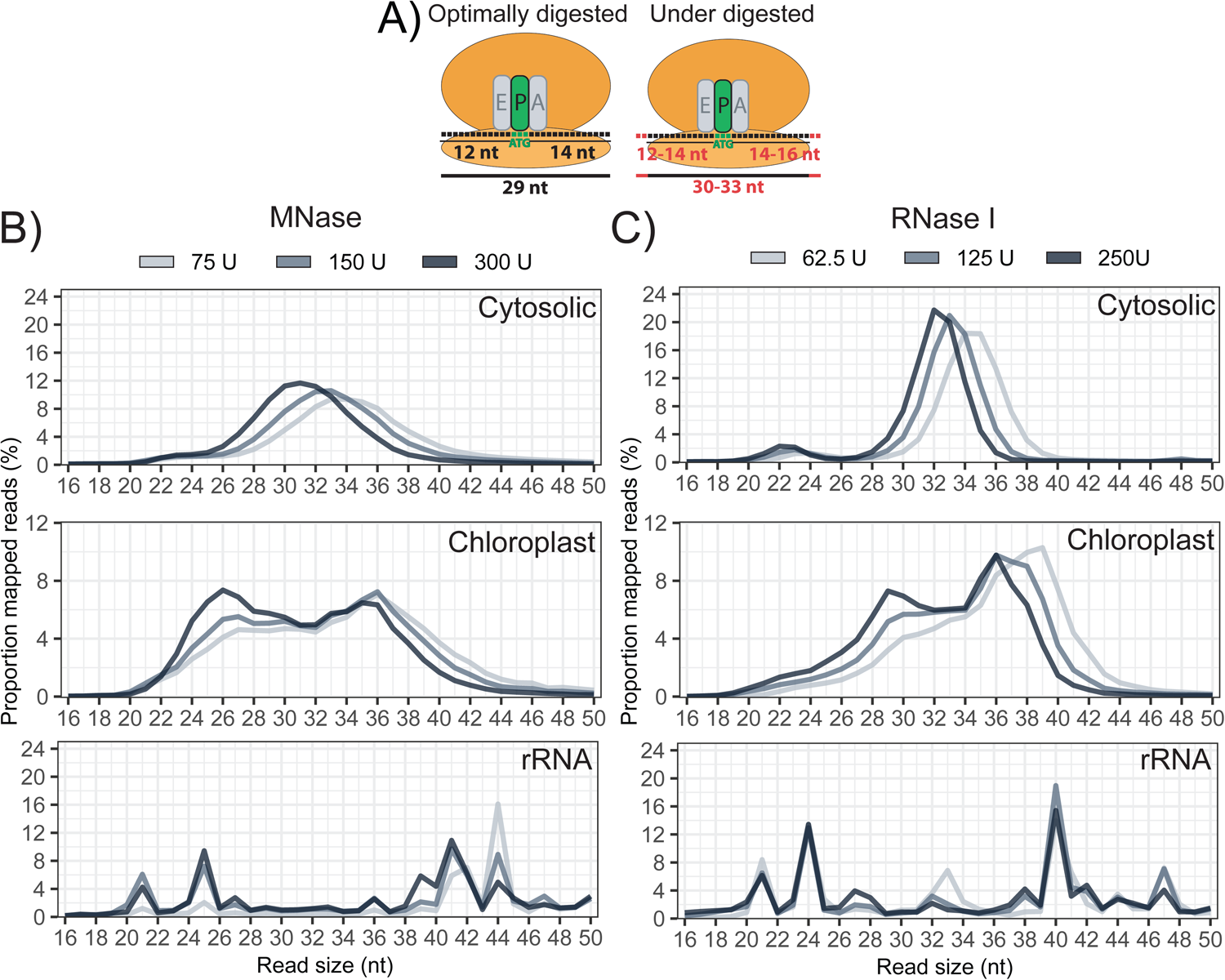
RPF size distribution across variable digestion conditions. **(A)** Schematic comparison between optimally digested and under-digested RPFs. Ribosomes decode the codon positioned in the ribosome P-site, which is typically 12 nt from the 5’ end from an optimally digested RPF. Exact P-site position is difficult to assess for under-digested RPFs, because of variable nucleotide digestion on the terminal ends (red). **(B-C)** Size distribution of RPFs mapping to the nuclear genome, the chloroplast genome, and rRNA, following digestion using (B) MNase or (C) RNase I.

Given that the size of cytosolic RPFs were larger than expected, additional increasing increments of RNase I (400 U, 550 U, and 700 U) were tested to find the minimum concentration that efficiently produces cytosolic RPFs sized 28-29 nt. These Ribo-seq libraries were remarkable similar in composition, counts over gene CDS, and triplet periodicity (Figure 5A-C), indicating that this digestion range very robustly generates reproducible data. Notable improvements in triplet periodicity were observed (55.4 – 59.9 % RPFs in frame 1, Figure 5C) compared to the under digested samples (43.1-49.1 % RPFs in frame 1, Figure 3B). When using 550-700 U of RNase I, the majority of cytosolic RPFs were stably around 29 nt (Figure 5D). Of note, an independent study following a similar ribosome profiling methodology used 1400 U of RNase I, and reported similar periodicity values and RPF size distribution around 29 nt (Chotewutmontri et al., 2018). This suggests that digestion beyond 700 U (at least up to 1400 U) does not provide cost effective benefits. Together, these observations prompted us to apply 600 U RNase I as standard procedure.

**Figure 5.**
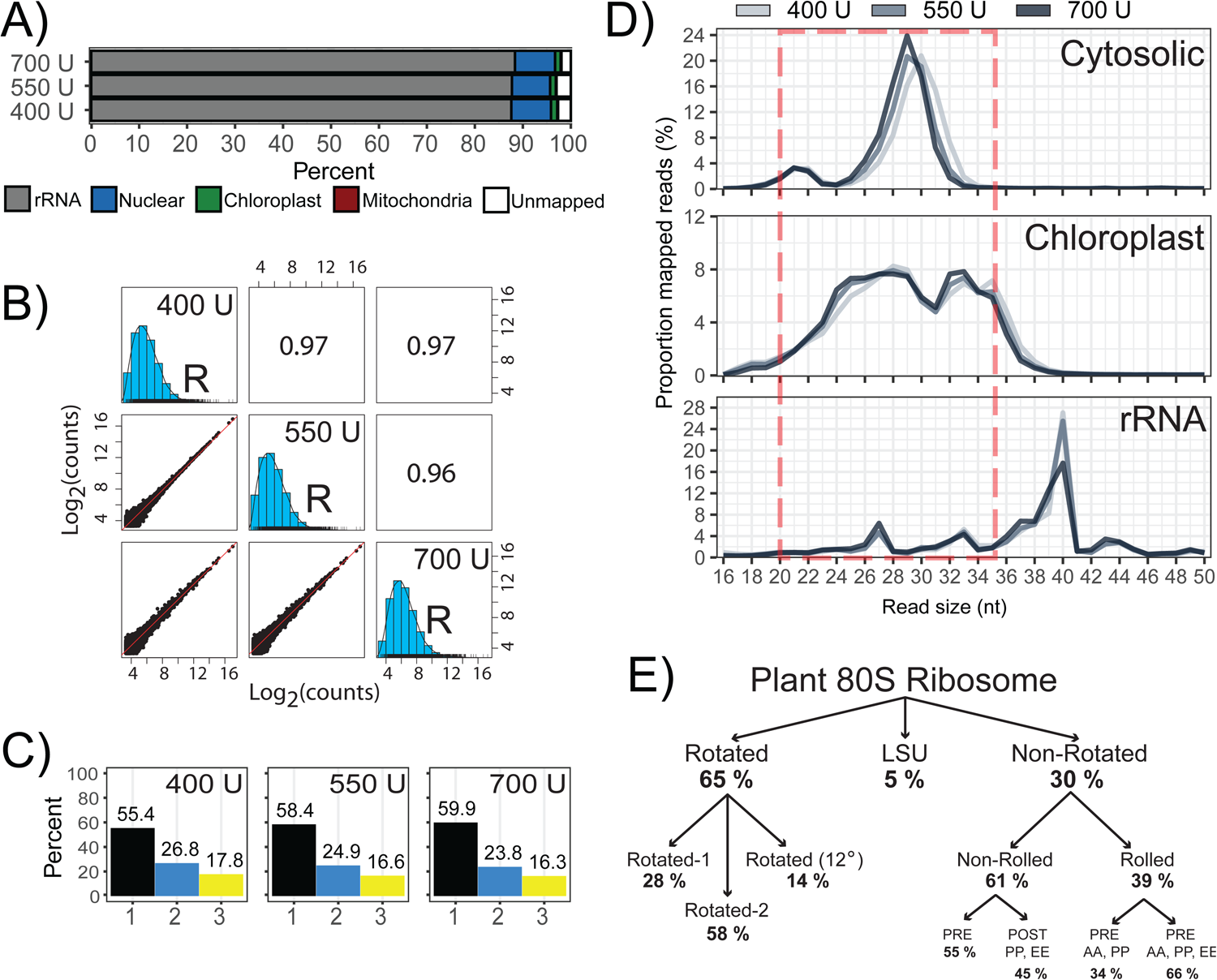
Refinement of RNase I treatment. Three additional Ribo-seq libraries were generated with increasing increments of RNase I treatment. **(A)** Ribo-seq library composition, as described in Figure 2. **(B)** Spearman’s correlation of RPF density over annotated CDS (log_2_-counts) as described in Figure 2. **(C)** Quantification of RPFs in the three frames of translation. Frame 1 is in reference to annotated start codons. **(D)** Size distribution of RPFs mapping to the nuclear genome, the chloroplast genome, and rRNA. The red dashed box indicates our new preferred RPF size selection to avoid major rRNA while including cytosolic and chloroplast RPFs. **(E)** Structural conformations of actively translating 80S cytosolic ribosomes from tobacco. The nomenclature and values were derived from CryoEM data from (Smirnova et al., 2023). LSU, Large Subunit of the ribosome. U, Units of applied ribonuclease; R, RNase I.

Actively translating 80S ribosomes undergo numerous conformational changes during an elongation cycle. In tobacco, a recent Cryo-EM study (Smirnova et al., 2023) has revealed that in a given snapshot, the majority of cytosolic ribosomes are found in the rotated (65%) and non-rotated (∼30%) states (Figure 5E). Although our triplet periodicity values are relatively low compared to other studies, we note that they stabilize at around 60%, which is similar to the proportion of ribosomes found in the rotated conformation. Since our protocol was optimized to minimize nuclease treatment, we speculate that the stabilized periodicity values around 60% are reflective of the diverse ribosome conformations. Although it is undeniable that higher periodicity can provide greater confidence in classifying actively translated ORFs, our experience with non-periodic datasets (generated using MNase) is that the observation of RPF density near the putative start and stop codon of an ORF is more than sufficient. In addition, the RPF distribution between non-periodic and tri-periodic datasets is highly similar, with the majority of reads mapping to CDS (Figure 3A). ORF detection programs often require a minimum number of reads along an ORF, which argues that increasing sequencing depth provides more benefits than either selecting only triperiodic reads or improving tri-periodicity (potentially introducing bias in the native distribution of different ribosome conformations; Fig. 5E). For these reasons, we focused our efforts into rRNA removal, which is the most cost-effective way to increase the number of informative reads (i.e., RPF coverage).

### Removal of contaminating rRNA fragments

Our initial Ribo-seq datasets were generated by selecting RPFs from 20-50 nt, to ensure that the majority of chloroplast RPFs were captured. We now recommend a size selection of 20-35 nt, which still captures the majority of chloroplast RPFs, while simultaneously excluding the very abundant rRNA fragments at ∼40 nt (Figure 5D). When broken down, the most problematic rRNA fragments belong to the nuclear-encoded 25S, 18S, and 5.8S rRNAs (Nu 25S, Nu 18S, Nu 5.8S) and the chloroplast-encoded 23S rRNA (Cp 23S), irrespective of nuclease treatment or plant material (Figure 6A). The sum of all other rRNA species contributed less than 2.5%, and therefore their contaminating effect is neglectable. Next, we identified high coverage rRNA regions that were repeatedly detected across several datasets, and designed biotinylated oligos to target these regions for removal (Figure S2). While designing the oligos, we noticed that the relative abundance of some rRNA fragments displayed high variation, even among technically similar replicates. For example, two fragments derived from the Nu 18S and the Cp 23S differed by 15% and 10% of the total library size, respectively, from two libraries that differed only by PCR (Figure S3B). Since PCR can have such a profound effect on the abundance of rRNA fragments, we reasoned that rRNA removal is most robust when performed prior to PCR amplification (ideally prior to any enzymatic step) because the molar ratios of oligos to the contaminants are best maintained. We thus formulated our initial Arabidopsis depletion cocktail (Version 1, Table S1) and performed rRNA depletion directly on the gel-purified RPFs. Following this procedure, we effectively reduced rRNA contamination from 85% to ∼25%, which corresponds to a 7-fold improvement in informative reads (Figure 6B). Examination of the rRNA-depleted dataset revealed that new rRNA fragments began to disproportionately dominate the library, prompting us to add five additional oligos to our depletion cocktail (Version 2, Table S1). Surprisingly, the extra oligos did not yield any benefits (Figure 6B), indicating that there is a limitation in the number of oligos that will result in noticeable improvements. Indeed, oligo cocktails containing 60 oligos report only a 50 % rRNA reduction (Chotewutmontri et al., 2018) which is less efficient than our 29 oligos. Since 2-5 rRNA fragments can account for more than 90% of a Ribo-seq library (Berg et al., 2020), a minimal cocktail containing only 5-10 oligos may already provide sufficient benefits for most applications. Next, the same procedure was applied to formulate a tobacco-specific rRNA depletion cocktail (Table S2), which was also very effective at reducing rRNA from ∼75% to ∼40 % (Figure 6C).

**Figure 6.**
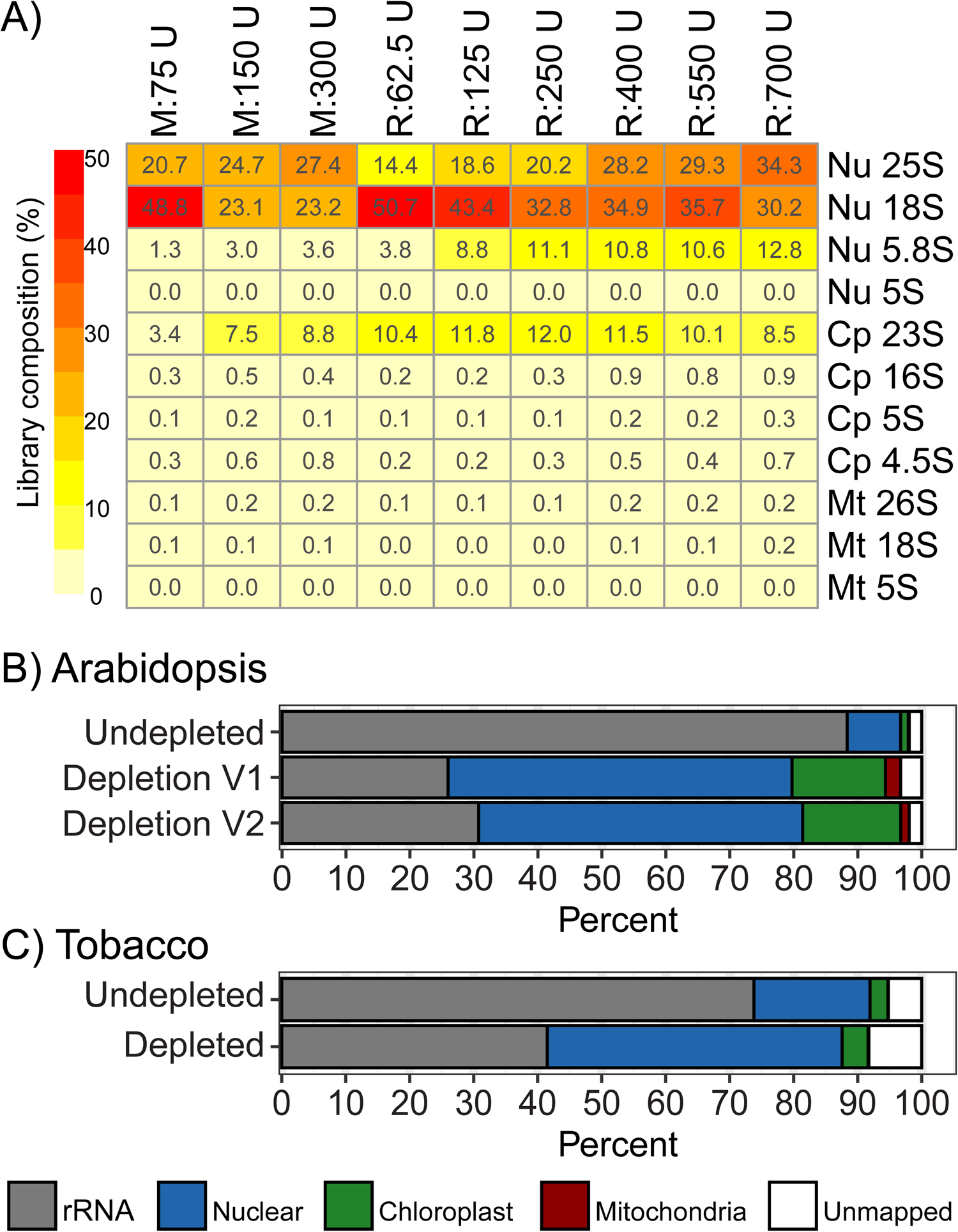
Removal of contaminating rRNA fragments from Ribo-seq datasets. **(A)** Contribution of individual rRNA species in Arabidopsis Ribo-seq libraries generated following different ribonuclease treatments. **(B)** Arabidopsis Ribo-seq library composition following rRNA depletion with depletion cocktail version 1 (V1) containing 24 oligos and depletion cocktail version 2 (V2) containing 29 oligos. The undepleted library corresponds to the 700 U RNase I-treated sample described above (Figure 5A). **(C)** Tobacco Ribo-seq library composition following rRNA depletion. The sequences of the biotinylated oligos used for rRNA depletion are provided in Supplemental Tables S1 and S2. Nu, nuclear-encoded; Cp, chloroplast-encoded; Mt, mitochondria-encoded; U, Units of applied ribonuclease; M, MNase-treated; R, RNase I-treated.

### Preservation of rRNA fragments and their spatial distribution within the 80S ribosome

As mentioned at the beginning, an initial concern was that increasing nuclease digestion would create more rRNA fragments. However, our data demonstrates that this is not the case over a wide range of nuclease concentrations. Despite the observation that RNase I concentrations higher than 250 U caused monosome breakdown (Figure 1C), we did not observe altered rRNA distributions (Figure 5D) or higher rRNA contamination (Figure 2A, Figure 5A) from treatments with higher nuclease concentrations, suggesting that no new fragments are formed. In fact, a closer examination revealed that the most abundant rRNA fragments are preserved across all our datasets, irrespective of the plant material or ribonuclease treatment (Figure 7A,B and Figure S2). Furthermore, many of the rRNA fragments identified in Arabidopsis are also present in tobacco Ribo-seq datasets (Figure 7C,D and Figure S4), suggesting that similar fragments are also preserved across plant species (i.e., in rRNAs orthologs).

**Figure 7.**
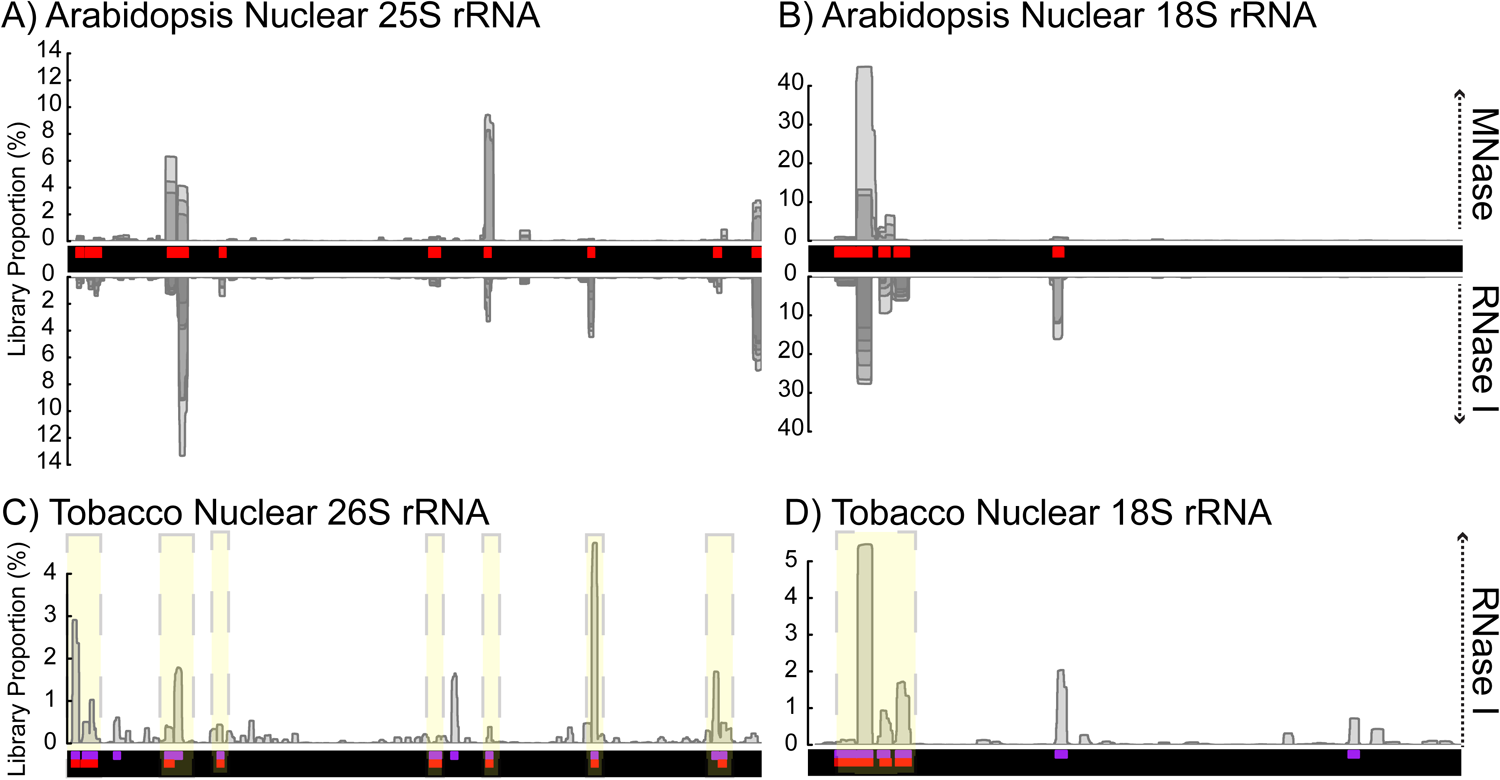
Preservation of contaminating 80S rRNA fragments. **(A-B)** Contaminating rRNA fragments under different ribonuclease regimes for the Arabidopsis nuclear-encoded (A) 25S and (B) 18S rRNAs. The coverage is derived from all Ribo-seq libraries described above that were generated with MNase (above y-axis) and RNase I (below y-axis). Regions with overlapping coverage from multiple libraries are shown in darker shades of grey. Arabidopsis depletion oligos (red) are given in Table S1. Similar plots for all rRNA species are shown in Figure S2. **(C-D)** Contaminating rRNA fragments across plant species, illustrated within the tobacco nuclear-encoded 26S (C) and 18S rRNAs (D). Depletion oligos used for tobacco (purple) are described in Table S2. For comparative purposes, the Arabidopsis depletion oligos (red) were aligned to tobacco rRNA genes, and are displayed on equivalent positions. Yellow shades indicate preserved rRNA contaminants between Arabidopsis and tobacco. Similar plots for all rRNA species are available in Figure S4.

Together, these observations prompted us to explore the spatial distribution of the most abundant rRNA fragments within the plant 80S ribosome to gain insights into their origin. To this end, we used the recently solved cryo-EM structure of the tobacco 80S ribosome (PDB: 8B2L)(Smirnova et al., 2023). The analysis confirmed that many of the rRNA fragments that regularly contaminate Ribo-seq datasets are derived from surface exposed rRNA helices which are not shielded by ribosomal proteins (Figure 8). The RNase nick sites occur more frequently on rRNA hairpins, loops, and bulges. Several of the most abundant contaminants (C1, C2, C10 and C14) are located on the solvent-exposed surface of the ribosome, which is where rRNA expansion segments (ESs) are predominantly localized (Yusupova and Yusupov, 2017). In contrast, few contaminants were localized at the subunit interface (contact site of the small and large subunits). The interface contains the three tRNA-binding sites (A, P, and E), the decoding center, and the peptidyl transferase center (Yusupova and Yusupov, 2017), and is well shielded from the environment. We reason that fragments corresponding to the interface and other well-protected regions of the ribosome are likely to be larger than 50 nt, and are thereby excluded following our applied RPF size selection (20-50 nt).

**Figure 8.**
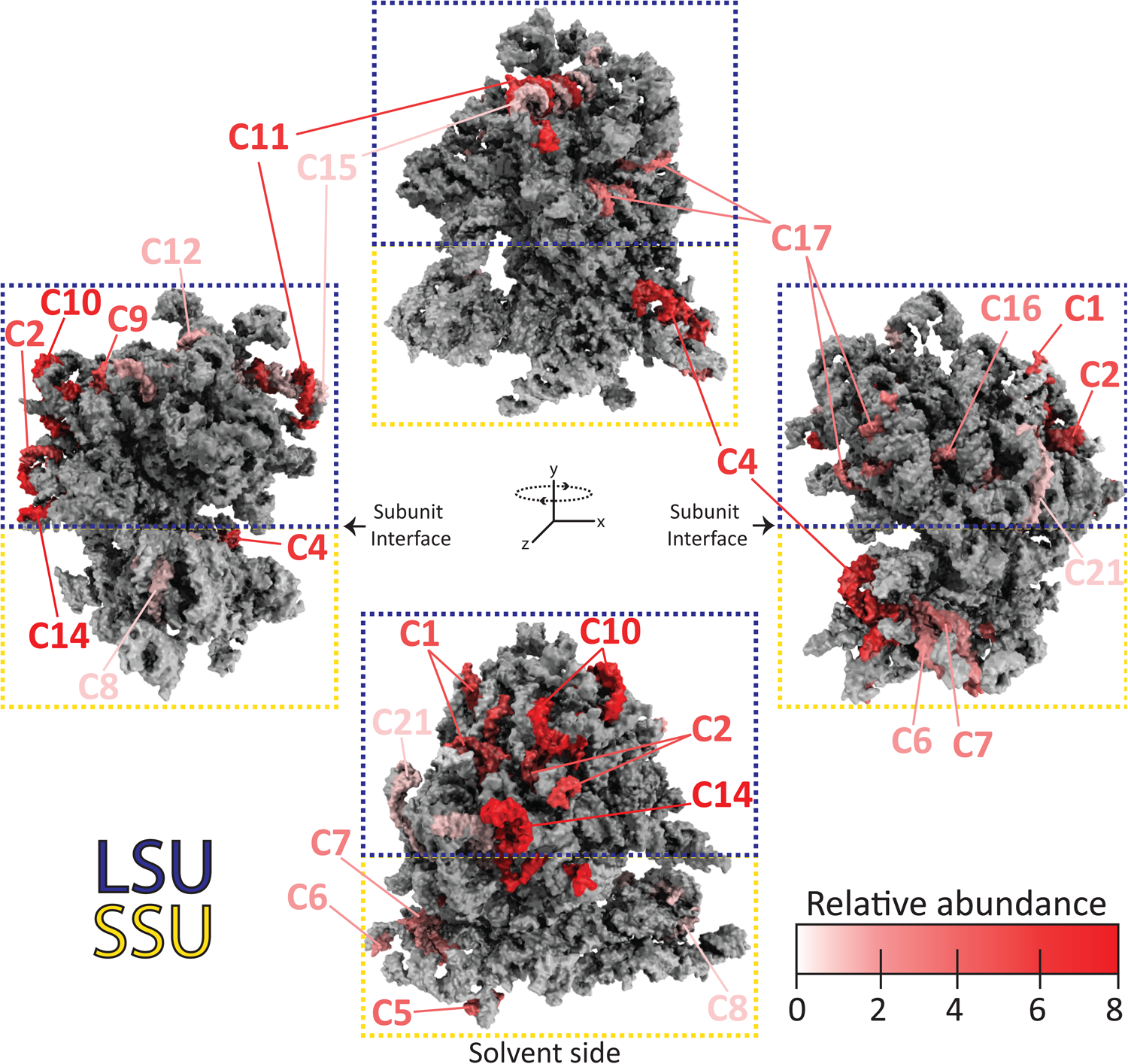
Mapping of contaminating rRNA fragments to the tobacco 80S ribosome structure. Major contaminating rRNA fragments (C1-C21) identified from tobacco Ribo-Seq datasets (Table S2) were mapped onto the molecular model of the tobacco 80S ribosome (PDB: 8B2L). The four rRNA molecules are shown in gray as spheres. Contaminants are colored in shades of red that reflect their relative abundance within our Ribo-seq datasets (additional details provided in Table S2). The ribosomal proteins are excluded from the model for clarity. The large subunit (LSU) and small subunit (SSU) are indicated in the blue and yellow dashed boxes, respectively.

These results highlight that many of the commercial rRNA depletion kits used for RNA-seq cannot perform well in Ribo-seq experiments. This is especially true for kits that contain a limited number of probes that target highly conserved rRNA sequences. Such probes are unlikely to correspond to the same fragments generated following nuclease treatment, and are not combined in optimal molar ratios. It is also worth noting that we have attempted using our Arabidopsis depletion oligos on tobacco samples, which was anticipated to be effective given the fragment similarities. However, only moderate depletion was achieved, which could be attributed to tobacco-specific single nucleotide polymorphisms (SNPs) that presumably hindered hybridization of the Arabidopsis oligos. Thus, a universal plant Ribo-seq depletion cocktail is unlikely to provide highly efficient rRNA removal across many plant species. Overall, these observations confirm the intuitive notion that major rRNA contaminants that dominate Ribo-seq datasets are formed from rRNA fragments whose 5’ and 3’ boundaries are readily accessible for ribonuclease attack. The most vulnerable regions belong to those located on the solvent-exposed surface of the ribosome. For the establishment of Ribo-seq in new plant species, these observations may facilitate the *in silico* prediction of major rRNA contaminants without any pioneer sequencing runs.

### Minimizing PCR bias

Ribo-seq protocols include a PCR amplification step, which can be a major source of bias when preparing sequencing libraries (Daniel et al., 2011). Indeed, we have observed that variation in PCR amplification can outweigh even differences in nuclease treatment (Figure S3A). To maximize reproducibility, libraries should be amplified to a similar concentration range using the same number of PCR cycles. In addition, libraries should not be amplified past the exponential phase of PCR, where the substrates of the PCR reaction become limiting and chimeric species begin to form. Although these notions seem trivial, we initially struggled with fulfilling both requirements because of highly variable PCR amplification across samples (12 – 19 cycles), despite using the same amount of RPF template. We suspect that this was caused by salts and/or pH altering molecules (or other contaminants) that co-purify with RPFs, and that negatively affect the enzymatic steps of the library preparation kit. This issue was alleviated by subjecting RPFs through an RNA purification column (e.g., NEB Monarch RNA cleanup) prior to library preparation, which has become a standard in our lab when using any library preparation kit. To ensure that libraries are amplified within the exponential phase of PCR, a qPCR approach was adopted to quantify the template prior to library amplification (see Supplemental Methods) as it has previously been described for Ribo-seq in non-plant species (McGlincy and Ingolia, 2017).

Thus far, all of our Ribo-seq datasets were prepared using RNA-ligase based strategies, which display only moderate PCR efficiency when using rRNA-depleted samples as input. An appealing alternative are ligation-free approaches which utilize the template-switching ability of selected reverse transcriptases. These strategies are tailored for samples with low RNA input, and have already been successfully applied for Ribo-seq in mammals (Hornstein et al., 2016). To compare these two approaches in plants, we generated Ribo-seq data using a ligation-based kit (NextFlex small RNA-seq V4) and a ligation-free kit (Diagenode D-Plex small RNA-seq). The ligation-free approach was magnitudes more efficient, requiring only 8 PCR cycles to obtain sufficient library quantities for sequencing, compared to the 16 PCR cycles for the ligation-based approach (Figure 9A). Expectedly, triplet periodicity was lower for the ligation-free approach, which is due to the inability to distinguish 3’-terminal adenosine nucleotides that were enzymatically added, from those 3’-terminal adenosine nucleotides that truly belong to RPFs. Despite the reduced periodicity, the quantitative translatomes were still highly comparable (Figure 9B). Thus, for general applications where codon-resolution is not required (to detect, e.g., rare ribosome frame-shifting events), we recommend the ligation-free approaches which are more efficient and convenient.

**Figure 9.**
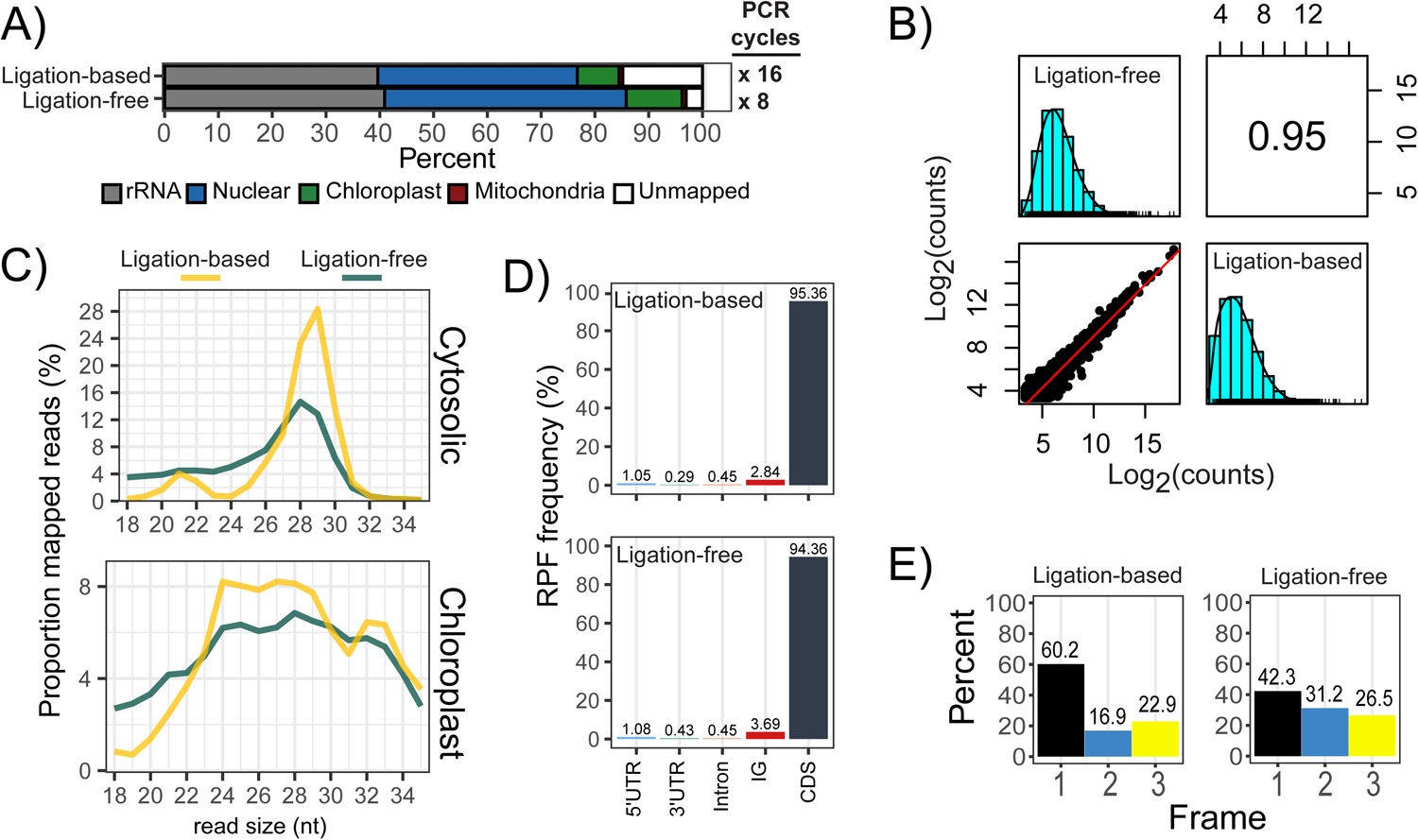
Comparison of ligation-based and ligation-free Ribo-seq strategies. Ligation-based strategies (e.g., NextFlex Small RNA-seq kit V4) use RNA-ligases for adapter attachment. Ligation-free strategies (e.g., Diagenode D-Plex small RNA-seq kit) use polyadenylation and reverse-transcription template switching for adapter incorporation. **(A)** Ribo-seq library composition. Indicated on the right are the number of PCR cycles required to obtain sufficient library quantities for sequencing. **(B)** Spearman’s correlation of RPF density over annotated CDS (log_2_-counts) as described in Figure 2. **(C)** Size distribution of cytosolic RPFs mapping to the nuclear genome, and chloroplast RPFs mapping to the chloroplast genome. **(D)** Proportion of mapped RPFs across genomic features. The percentages are calculated after removal of rRNA. **(E)** Quantification of RPFs in the three frames of translation. Frame 1 is in reference to annotated start codons.

### Analysis of Chloroplast RPFs

Plants harbor three translationally active compartments: the cytosol, mitochondria and plastids (predominantly chloroplasts in green tissue). While cytosolic RPFs clearly dominate plant Ribo-seq libraries and mitochondrial RPFs are neglectable, chloroplast RPFs make up a substantial fraction (Figure 6). Due to the fact that essential proteins of the photosynthesis machinery are chloroplast-encoded, chloroplast translation is essential to establish photosynthesis. For studies that focus solely on chloroplast translation, we find that library sizes of 2-5 Million reads (after rRNA depletion) provide sufficient coverage for the vast majority of chloroplast genes. Due to the structural differences between the eukaryotic 80S ribosome of the cytosol, and the prokaryotic-like 70S ribosome of the plastid, it is recommended that P-site offsets are estimated separately for these two ribosome species. The P-site offsets for cytosolic RPFs are predominantly 12-13 nt from the 5’end (Figure S5), which is the norm for eukaryotic RPFs (Lauria et al., 2018). In contrast, the P-site offsets for the chloroplast RPFs are diverse, and decrease from the 5’end as the RPF gets smaller (Figure 10A). This indicates preferential nuclease digestion from the 5’end, which is a pattern that has previously been observed for chloroplast RPFs (Chotewutmontri and Barkan, 2016). It was reported before that the determination of chloroplast P-site offsets can be performed by applying a constant 7 nt from the 3’end (Chotewutmontri and Barkan, 2016). When applying 3’mapping to our own dataset, similar offset values (6-8 nt) were only observed for smaller RPFs (20-30 nt), whereas the larger RPFs (31-40 nt) displayed a constant 15 nt offset (Figure 10A). It should be noted that chloroplast metagene analyses are inherently noisier since most land plant chloroplast genomes only encode ∼80 CDS genes. Furthermore, some chloroplast transcripts are polycistronic with very short spacers in between reading frames (or even overlapping reading frames), thereby making it difficult (or impossible) to distinguish terminating ribosomes from initiating ribosomes around these short spacers. Despite these limitations, triplet periodicity is still visible across chloroplast genes (Figure 10B,C).

**Figure 10.**
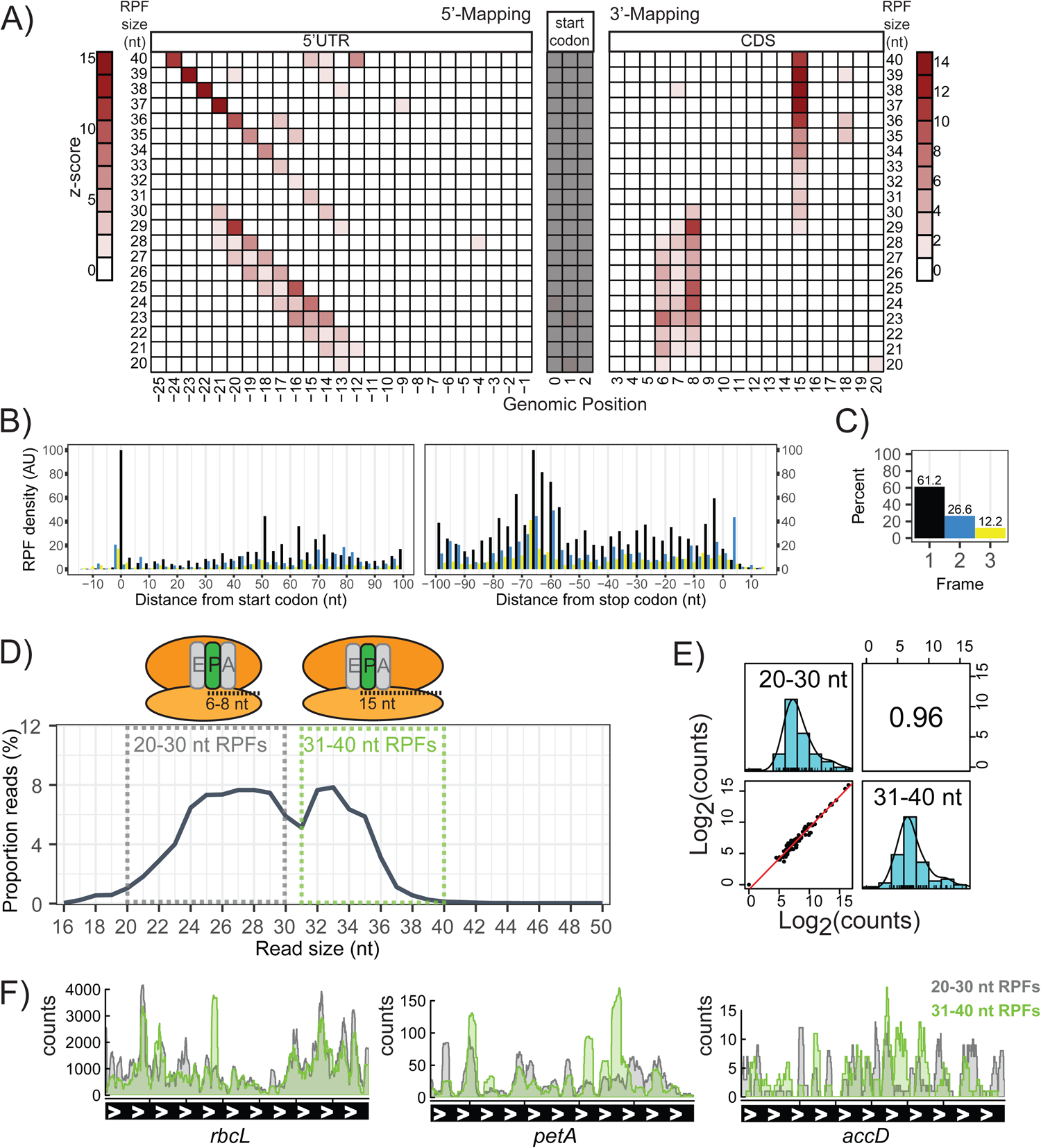
Properties of chloroplast RPFs. **(A)** P-site offsets. The RPF density around the start codon is normalized for each RPF size. Densities for 5’-mapped (left) and 3’-mapped (right) are both shown. **(B)** Metagene analysis for RPFs over the 79 annotated CDS genes of the chloroplast. RPFs are positioned at their P-site using the 5’-mapping offsets determined in (A). **(C)** Quantification of RPFs in each frame of translation. Frame 1 is in reference to the start codon. **(D)** Bimodal RPF size distribution corresponds to unique P-site offsets. The small (20-30 nt) and large (31-40 nt) RPFs are indicated in dashed grey and green boxes, respectively. **(E)** Spearman’s correlation of RPF density over chloroplast CDS (log_2_-counts), for the small- and large-sized RPFs. **(F)** Coverage of the small- and large-sized RPFs over chloroplast genes. Highly expressed (*rbcL*), medium expressed (*petA*), and lowly expressed (*accD*) genes were selected to provide a broad representation. The data is derived from the library corresponding to the 700 U RNase I treatment (described in Figure 5).

Interestingly, the small and large RPFs that are characterized with the distinct P-site offsets, correspond to the two visible peaks in the RPF size distribution (Figure 10D). To explore this further, chloroplast RPFs were size separated *in silico*, to determine if the small and large RPFs display unique localization patterns. Both RPF populations were similarly distributed across all chloroplast genes indicating no bias towards specific genes (Figure 10E). For eukaryotic ribosomes, a smaller size (∼19-21 nt) corresponds to stalled ribosomes containing an empty A-site (Wu *et al*., 2019a). This is unlikely to be the case for the small RPFs of the chloroplast, since they are relatively abundant (Figure 10D) and are widespread along the entire CDS (Figure 10F). Hence the molecular cause for the two observed RPF sizes remains to be determined. It is tempting to speculate that the small and large RPF populations of the chloroplast represent different rotational conformation states of actively translating ribosomes.

### Complementary transcriptome

For calculating translation efficiency (TE), complementary RNA-seq libraries are typically generated in parallel with Ribo-seq libraries. Since the Ribo-seq dataset described here were generated from optimization trials, complementary transcriptomes were not generated, so no TE calculations are provided. However, we want to share our experiences with transcriptome generation: The depletion of rRNA from total RNA is standard in RNA-seq, with the most popular methods being enrichment of polyadenylated (poly(A)) transcripts, subtractive hybridization with biotinylated oligos, and enzymatic digestion. Since chloroplast RPFs contribute substantially to the plant translatome, we prefer strategies that preserve chloroplast transcripts, which is why we avoid poly(A) mRNA enrichment (since chloroplast transcripts are regularly not polyadenylated). We have also tested commercial rRNA removal kits that utilize subtractive hybridization, and have had good experience from oligos derived from riboPOOLs (siTOOLs Biotech). However, we observed a new problem that arises following efficient rRNA removal. Due to PCR-biases, other abundant RNA species sometimes disproportionately dominate a sequencing libraries. For this reason, we prefer to use enzymatic based depletion strategies (e.g., Zymo-Seq RiboFree Total RNA Library Kit) which remove abundant RNA species in a sequence-independent manner. This strategy removes most rRNA, preserves organelle transcripts, and prevents any single RNA species from becoming disproportionately over represented.

## Conclusions

The genome-wide analysis of translation was revolutionized by ribosome profiling, which is often optimized across different labs to suite individual purposes. Through our own optimization efforts for Arabidopsis and tobacco, the methodology described here focuses on minimizing nuclease treatment and preserving chloroplast RPFs. This necessitates a broader RPF size selection, which comes at the expense of lower triplet periodicity. However, it has been demonstrated that non-periodic data (generated from MNase) still provides accurate translational dynamics (Zoschke and Barkan, 2015; Gawroński et al., 2018; Trösch et al., 2018; Schuster et al., 2020; Gao et al., 2022). Therefore, we instead prioritize sequencing depth, which we believe to be the limiting factor when trying to identify lowly translated ORFs. For this reason, we emphasize rRNA removal, which we find to be very efficient when performed at the RNA level, prior to any enzymatic steps. In addition, our structural assessment of rRNA fragments provide new insights that should benefit the general community when establishing ribosome profiling in new plant species. Together with our ribosome profiling protocol for the green alga *Chlamydomonas reinhardtii* (Gotsmann et al., 2023), this provides a tool box that paves the way for highly comparative Ribo-seq studies in a wide range of plant species.

## Supporting information

FileS1. Example scripts

## Acknowledgements

We thank Ines Gerlach (Max Planck Institute of Molecular Plant Physiology) for excellent technical assistance. We thank Alice Barkan and Prakitchai Chotewutmontri (University of Oregon) for helpful discussions at the early stages of the optimization process of our Ribo-seq protocol. We acknowledge the excellent sequencing service of the Sequencing Core Facility of the Max Planck Institute for Molecular Genetics.

## Author Contributions

MKYT, RB, YG, RG and JHL generated the Ribo-seq libraries. MKYT, VG, and AF provided bioinformatics analysis. FMS and JS performed the mapping of the rRNA fragments onto the ribosome structure. YG performed the polysome analysis. MKYT and RZ wrote the manuscript, with contributions from FW and MJH.

## Declarations Ethical approval

Not applicable

## Availability of data and materials

The sequencing data have been deposited in NCBI’s Gene Expression Omnibus under accession number GSE226508.

## Supplemental Data

**File S1**. Example scripts

**Table S1.**
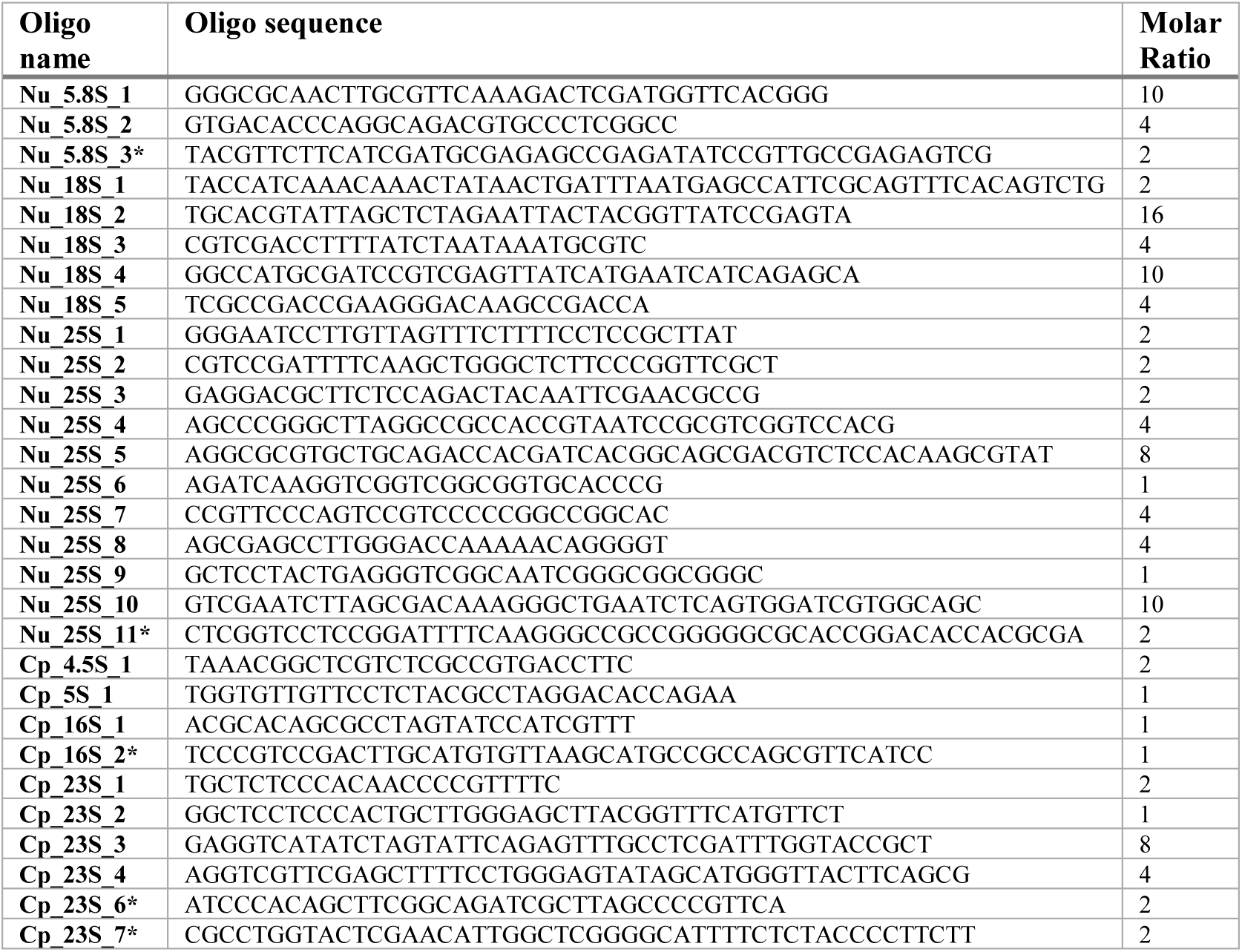
Custom biotinylated DNA depletion-oligos for rRNA removal from Arabidopsis RPF samples. Depletion oligos were designed for Arabidopsis rRNA loci with high coverage across all Arabidopsis Ribo-seq libraries produced from our lab (for details see Results). The molar ratio reflects the relative abundance of these contaminants among all our Ribo-seq libraries. The final oligo cocktail (version 2) contains all the oligos in this table, mixed at the indicated molar ratios. The preliminary oligo cocktail (version 1) contained only the oligos without an Asterisk (*). Nu, nuclear-encoded; Cp, chloroplast-encoded.

**Table S2.**
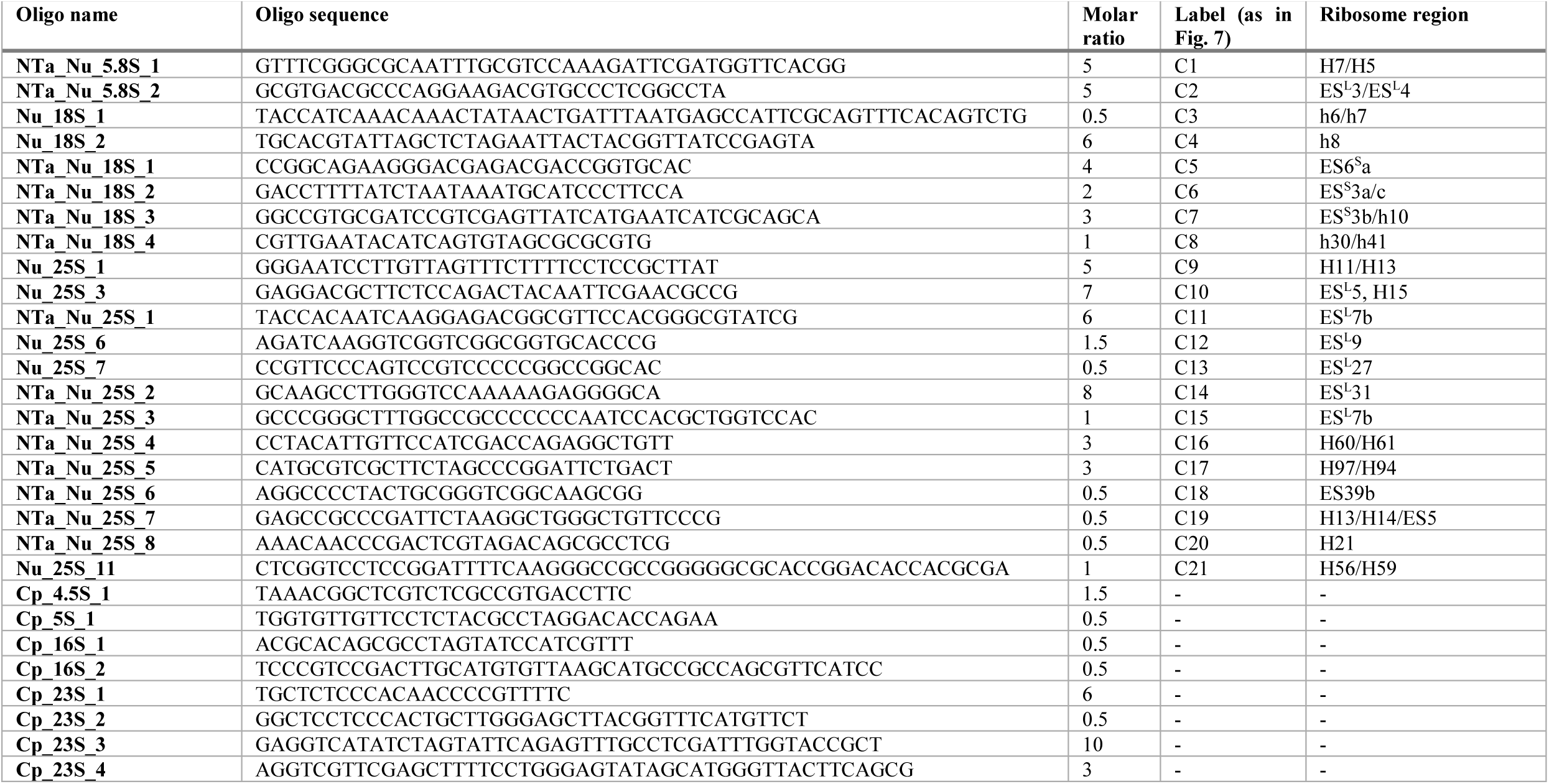
Custom biotinylated DNA depletion-oligos for rRNA removal from tobacco RPF samples. Depletion oligos were designed for tobacco rRNA loci with high coverage across tobacco Ribo-seq libraries produced from our lab (includes datasets not described in this study). The molar ratio reflects the relative abundance of these rRNA fragments across our Ribo-seq libraries. The “Labels” of the rRNA fragments depicted in Figure 7, are listed accordingly. The “Ribosome regions” are named accordingly to Armache et al, 2010. Oligos prefixed “NTa” are used only for tobacco rRNA depletion, whereas those without prefix are used for both Arabidopsis and tobacco rRNA depletion. Nu, nuclear-encoded; Cp, chloroplast-encoded; LSU, large subunit; SSU, small subunit.

### Optimized Plant Ribosome Profiling protocol

#### Isolation of ribosome-protected fragments and total RNA

300 mg frozen plant tissue was homogenized in liquid nitrogen with a mortar and pestle followed by the addition of 3 mL ribosome extraction buffer (0.2 M sucrose, 0.2 M KCl, 40 mM Tris-OAc pH 8.0, 10 mM MgCl_2_, 10 mM 2-Mercaptoethanol, 2% (v/v) polyoxyethylene (10) tridecyl ether, 1% (v/v) Triton X-100, 100 μg/mL chloramphenicol, 100 μg/mL cycloheximide). After brief mixing, a 0.5 mL aliquot of the lysate was flash frozen and stored at −80 °C for later total RNA extraction using TRIzol reagent (ThermoFisher cat# 15596026). This total RNA can be used for standard RNA-seq experiments. This step is especially important if the calculation of translation efficiencies is desired (i.e., the normalization of Ribo-seq data to RNA-seq data). The remaining lysate was filtered through glass wool, followed by centrifugation for 10 min at 15,000 x *g* at 4 °C to remove cell debris. 2.5 mL of the clarified lysate were incubated with 1500 U of RNase I (Ambion cat# AM2294) for 1 h at room temperature with gentle rotation to degrade mRNA regions that are not covered and protected by translating ribosomes. The ribonuclease-treated lysate contains mainly monosomes which were loaded onto a 2 mL sucrose cushion (30% (w/v) sucrose, 40 mM Tris-Acetate pH 8.0, 100 mM KCl, 15 mM MgCl_2_, 0.1 mg/mL chloramphenicol, 0.1 mg/mL cycloheximide, 0.2% β-MeEtOH) and centrifuged (OptimaTM L-80 XP Ultracentrifuge - Beckman Coulter, SW55 Ti rotor) for 1.5 h at 303,800 x *g* at 4 °C. The supernatant was then carefully aspirated and the pelleted monosomes resuspended in 0.5 mL footprint isolation buffer (10 mM Tris pH 8.0, 1 mM EDTA pH 8.0, 100 mM NaCl, 1% (w/v) SDS, 0.1 M EGTA pH 8.0). The RNA was immediately extracted from this pellet using 0.5 mL TRIzol reagent. Next RPFs were size-selected through electrophoresis on a 12% denaturing polyacrylamide gel (19:1, acrylamide:bisacrylamide) prepared in 1x TBE buffer (89 mM Tris, 89 mM Boric Acid, 2mM EDTA pH 8.0) containing 8 M urea. To this end, 25 μg RNA were resuspended in 40 μL of ribosome footprint loading buffer (90% (v/v) deionized formamide, 20 mM Tris-HCl pH 7.5, 20 mM EDTA pH 8.0, 0.04% (w/v) bromophenol blue, and 0.04% (w/v) xylene cyanol) and denatured for 10 min at 70 °C. The gel was run in 1x TBE buffer with a constant power of 30 W at constant temperature of 12 °C (achieved by a cooling unit). Co-migrating pre-stained RNA ladder (Biodynamics Laboratory cat# DM253) was used to visualize the regions of the gel to excise RPFs (20-35 nt). RNA was eluted from the excised gel piece in 4 mL TESS (10 mM Tris pH 8.0, 1 mM EDTA pH 8.0, 0.1 M NaCl, 0.2% (w/v) SDS) by overnight incubation at 4 °C with gentle rotation. Eluted RNA was isolated with 4 mL of phenol:chloroform:isoamyl alcohol (25:24:1), followed by overnight ethanol precipitation at −20 °C. To increase purity and further narrow the volume, the RNA pellet was resuspended in 0.1 M NaCl (500 μL) and subjected to a second round of phenol:chloroform:isoamyl alcohol (25:24:1) extraction, followed by two washes with chloroform:isoamylalcohol (24:1) and overnight ethanol precipitation at −20 °C. The received RPF pellet was washed twice with 75% ethanol and resuspended in 20 μL RNase-free water. Typical yields are 200-600 ng of RPFs per 300 mg plant fresh weight (depending on tissue source, e.g., developmental stage, mutant phenotype, etc.).

#### rRNA depletion

RNase free Biotinylated oligos were purchased from metabion international (Planegg, Germany). Oligo hybridization was performed following an adapted protocol from (Kraus et al., 2019). 150 ng of gel-purified RPFs were mixed with the following components in a PCR tube: 4 μL deionized formamide, 1 μL 20X SSC (3M NaCl, 0.3M Na-Citrate, pH 7.0 with HCl), 2 μL EDTA (5 mM, pH 8.0), 0.7 μL biotinylated oligo mix (100 µM), and RNase free water up to 20 μL. Hybridization of the oligos was performed in a thermocycler with heated lid using a slow temperature ramp according to the table below:

|  |  |  |  |  |  |  |  |  |  |  |
| --- | --- | --- | --- | --- | --- | --- | --- | --- | --- | --- |
| Temp ($^{\circ}$C) | 80 | 75 | 70 | 65 | 60 | 55 | 50 | 45 | 40 | 35 |
| Time (min) | 5 | 2 | 2 | 2 | 2 | 2 | 2 | 2 | 2 | 2 |
| Ramp rate | -0.1$^{\circ}$C/s | | | | | | | | | |

After oligo hybridization, each sample was topped off to 40 µL using a solution of 1X SCC and 20% formamide and kept at 35 °C until ready for oligo removal, which was performed using Dynabeads MyOne C1 (Thermo Fisher cat# 65002). For each sample, 45 µL of beads were washed according to manufacturer’s protocol for RNA applications, and divided into 30 µL and 15 µL aliquots for performing two rounds of oligo removal. Each sample was incubated with the 30 µL bead aliquot at room temperature for 15 min, and magnetized for 2 min (the supernatant contains the partially rRNA-depleted sample). The second round of oligo removal was performed by aspirating the supernatant into the 15 µL bead aliquot and repeating the incubation and magnetization. The final supernatant contains the rRNA-depleted RPFs, which was aspirated, transferred into a clean tube and ethanol precipitated overnight at −20 °C. The rRNA-depleted RPF pellet was resuspended in 42 µL RNase-free water and treated with TURBO DNase (Thermo cat# AM2238) in a 50 µL reaction according to the manufacturer’s instructions, in order to remove traces of leftover DNA oligos. Following DNase treatment, RPFs were purified using the Monarch RNA cleanup kit (NEB cat# T2030) following the modified protocol for small RNA, and eluted in 35 µL RNase-free water. Typical RPF yields following depletion are ∼30% (i.e., 50 ng rRNA-depleted RPFs from 150 ng gel-purified undepleted RPFs).

#### RPF Library preparation

For ligation-free Ribo-seq, rRNA depleted RPFs were directly used as input into the D-plex small RNA-seq kit (Diagenode cat#C05030001) according to manufacturer’s instructions, and amplified with 7-9 PCR cycles. For RNA-ligase Ribo-seq, rRNA-depleted RPFs were treated with T4 polynucleotide kinase (PNK; ThermoFisher, cat#EK0031) in a two-step reaction, to prepare the terminal ends for adapter ligation. First, the depleted RPFs (∼50 ng in 35 µL) were added to a 50 µL PNK reaction without ATP, for 10 min at 37 °C to promote the dephosphorylation reaction (removal of 3’ phosphate groups and 2’, 3’ cyclic monophosphate, which are typical end products of MNase and RNase I digestion). Next, ATP was added to the reaction, followed by another incubation for 30 min at 37 °C. The end-repaired RPFs were purified using the Monarch RNA cleanup kit using the modified protocol for small RNA, and eluted in 12 µL RNase-free water. These purified RPFs were immediately used as input for the NEXTflex small RNA-seq kit v3 (Perkin Elmer, cat# NOVA-5132-06) or v4 (Perkin Elmer, cat#NOVA-5132-31), according to the manufacturer’s instructions. The resulting cDNA was stored at −80 °C until the number of PCR cycles required for library amplification, was experimentally determined by qPCR (see below). In the final PCR amplification step of NEXTflex library preparation, the cycle number was adjusted according to the qPCR results, and typically require 14-16 PCR cycles.

#### qPCR to determine the cycle number for Ribo-seq library PCR amplification

From our experience, NEXTflex libraries that are <1 ng/µL are very difficult to multiplex in equimolar ratios, due to less accurate qubit quantification. In contrast, libraries >10 ng/µL can be considered to be overamplified and are outside the exponential PCR doubling range, creating technical biases in the results. The following qPCR protocol was designed to produce libraries within the range of 1.5-6.0 ng/µL. The primers are derived from the NEXTflex small RNA-seq kit v3:

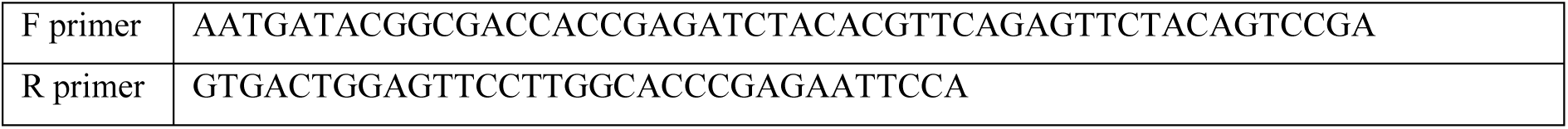

The instructions of the NEXTflex kit were followed up to STEP-F, eluting the cDNA product in 21 uL of elution buffer. Subsequently, 1 µL of this purified cDNA product was combined with 10 µL 2X Power SYBR Green PCR master mix (catalog# 4368577), 2 µL forward primer (0.5 µM), 2 µL reverse primer (0.5 µM), and 5 µL water (total volume of 20 µL). This qPCR reaction is aliquoted into 3 technical replicates (5 µL reactions per well), and run on an Applied biosystems 7900 HT machine with the following settings:

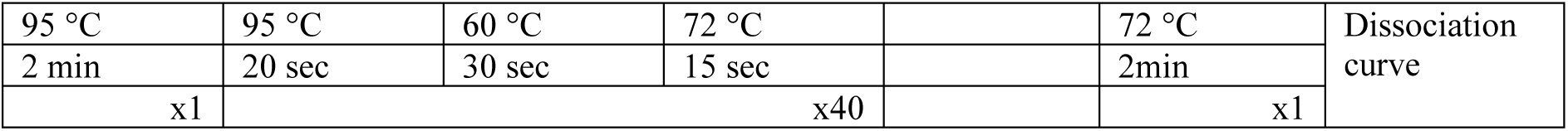

Based on the absolute Ct values obtained in this qPCR, the remaining 20 µL of the cDNA were PCR-amplified according to the table below:

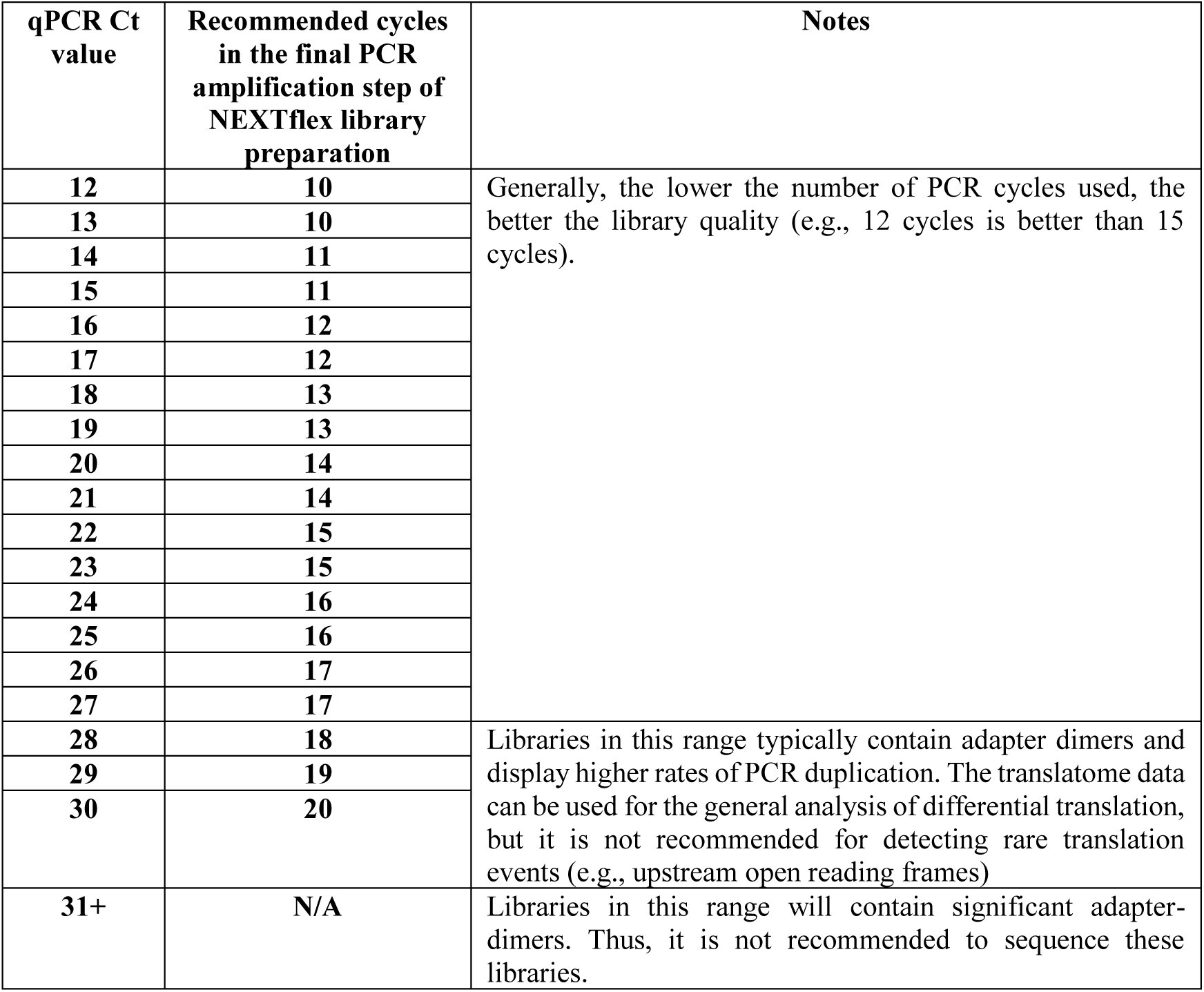

Absolute Ct values may vary across machines and applied qPCR kits. Hence, it is recommended to calibrate these values to your machine and kits, according to the final library concentration in pioneer experiments.

#### RNA seq library preparation

Although we do not include RNA-seq libraries in this manuscript, this is important to include in many ribosome profiling experimental setups to calculate translation efficiencies (as RPF abundancies normalized to transcript abundancies). In green plant tissue, up to a quarter of the RPFs are of chloroplast origin, so we prefer to use RNA-seq strategies that also preserve chloroplast transcripts. We routinely use the Zymo-Seq RiboFree total RNA library kit (Zymo cat# R3000/R3003) whose protocol depends on an enzymatic degradation of rRNAs.

#### Bioinformatics analysis

##### Arabidopsis thaliana

The Arabidopsis GTF gene annotations (v45) and TAIR10 genome were downloaded from ensembl (https://plants.ensembl.org/index.html). Cutadapt (Martin, 2011) was used to remove adapters and UMIs. Ribo-seq contaminants are generally defined as rRNA, tRNA and snoRNA. These sequences were extracted from the gene annotation file, and converted to fasta format using bedtools (Quinlan and Hall, 2010). Importantly, the rRNA gene annotations from TAIR10 (AT2G01010, AT2G01020, AT3G41768, AT3G41979, ATMG00020, ATMG01380, ATMG01390, ATCG00920, ATCG00950, ATCG00960, ATCG00970, ATCG01160, ATCG01170, ATCG01180, and ATCG01210) do not include the nuclear 25S and 5S. Thus, these sequences were obtained from NCBI (25S rRNA; X52320.1) and the 5S rRNA database (5S; http://combio.pl/rrna/). Contaminants were filtered out using STAR aligner (Dobin et al., 2013) with the following parameters: *--outFilterMismatchNoverLmax 0.1 --outReadsUnmapped Fastx –outSAMmultNmax 1*. In the datasets described here, tRNA and snoRNA contribute negligible amounts (<0.5%). Hence, “contaminants” will be synonymously used to describe rRNA. After contaminant filtering, the remaining reads were mapped to the genome using STAR aligner using the following parameters: *--outFilterMismatchNoverLmax 0.1 --alignIntronMax 8000 –outSAMmultNmax 1*. Coverage profiles along the genome, or rRNA were generated using bedtools and plotted using the Sushi package in R (Phanstiel, 2022). Mapped reads were counted over annotated CDS using FeatureCounts (Liao et al., 2014) using the parameters: *-M -s 1 -t CDS -g gene_id*. TMM normalization was performed using edgeR (Robinson et al., 2009), and genes with fewer than 10 counts were removed. P-site estimation, metagene analysis, and phase analysis were performed using the “plastid” package (Dunn and Weissman, 2016), and plotted in R. In order to quantify RPF density in each of the genomic features (CDS, 5’UTR, 3’UTR, introns, and intergenic regions), the Arabidopsis GTF file was processed using the “cs generate” function of “plastid” (Dunn and Weissman, 2016) to collapse multiple transcript isoforms into a single transcript isoform. This merged annotation file was used to create individual BED files that describe each of the genomic features. Only uniquely mapped RPFs were used for quantification within genomic features, which were extracted from the BAM alignment using samtools (Li et al., 2009). The genomic position and density of all RPF P-sites was calculated using the “make_wiggle” function of “plastid”. Quantification of RPF P-sites that overlap with each of the genomic features, was performed using the “map” function of “bedtools”.

##### Nicotiana Tabacum

There are two available genomes for tobacco, which are both available from solgenomics (https://solgenomics.net/). The most recent version was used for alignment (Edwards et al., 2017) since it is conveniently assembled into chromosomes. However, the Edwards genome assembly does not have annotations for tRNA. Therefore, tRNA sequences were obtained from the Sierro genome annotations (Sierro et al., 2014). The tobacco chloroplast (NC_001879.2) and mitochondrial (NC_006581.1) genomes were obtained from NCBI. Chloroplast and mitochondrial rRNA sequences were extracted from the respective genome annotations. Nuclear rRNA sequences were obtained from NCBI: 26S (AF479172.1), 18S (AJ236016.1), 5.8S (AJ300215.1) and 5S (AJ222659.1). The remainder of the analysis pipeline follows the same structure as what was performed for Arabidopsis. Additional parameters with example scripts can be found in Supplemental file 1.

