## Supplementary figures and images for "Guidelines for Performing Ribosome Profiling in Plants Including Structural Analysis of rRNA Fragments"

### Supplemental Figures

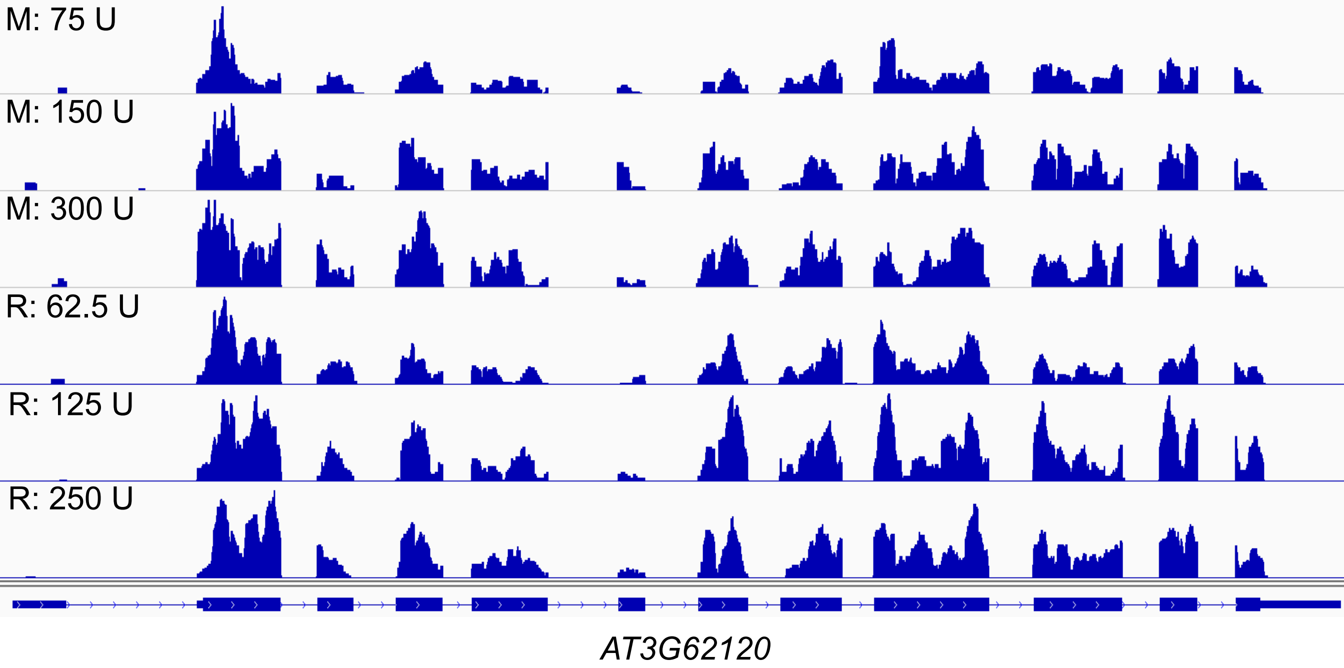


Figure S1

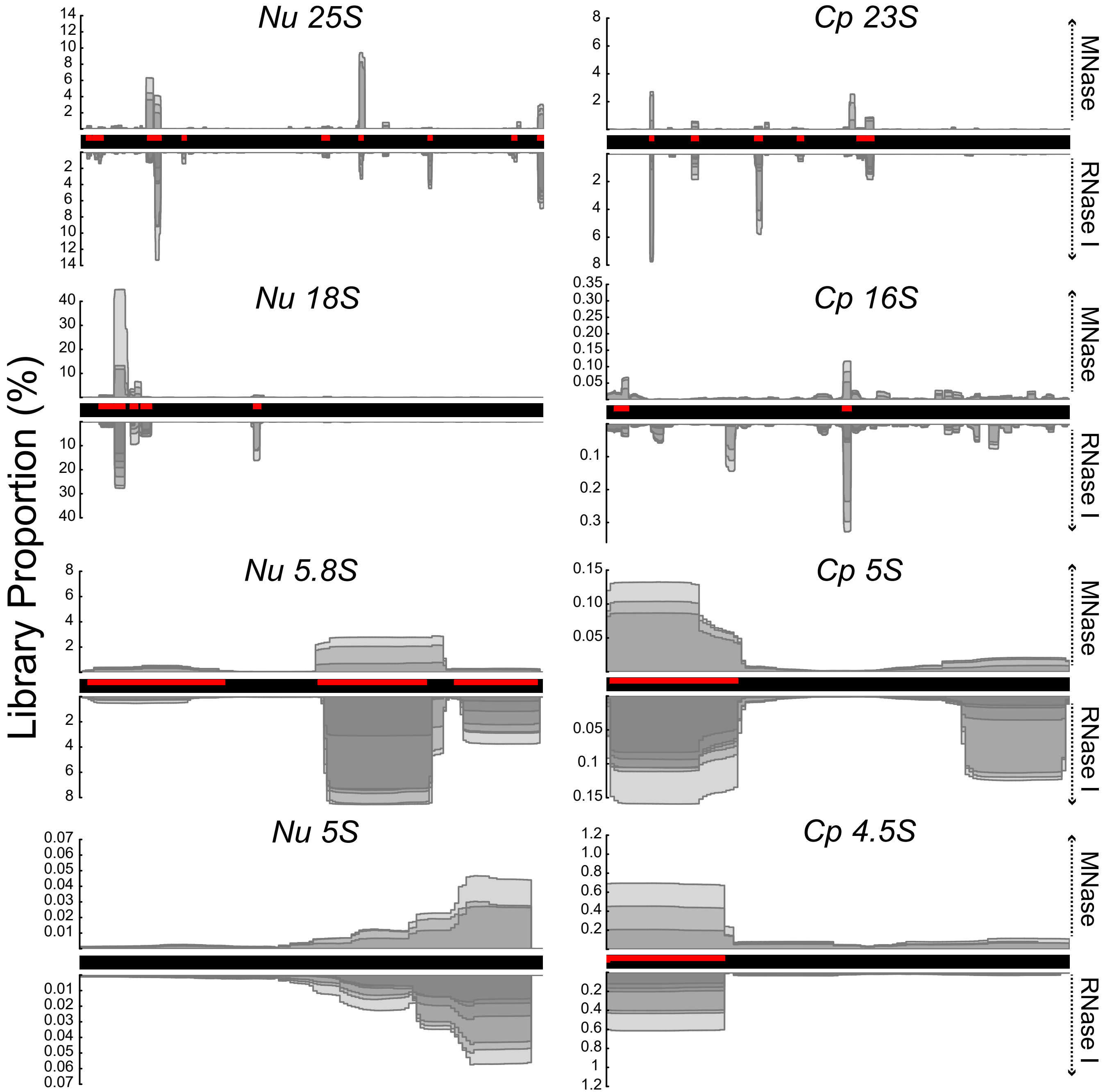

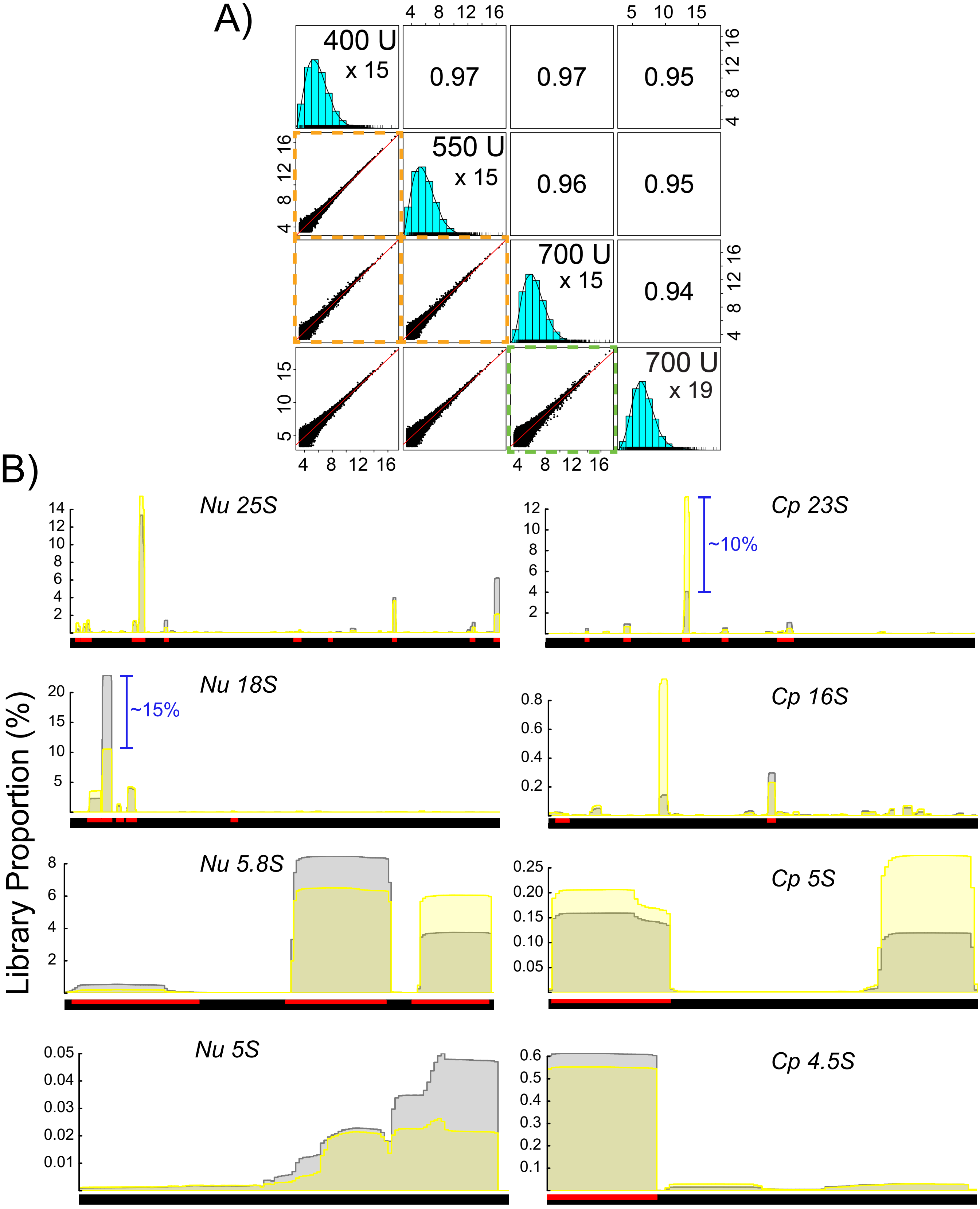

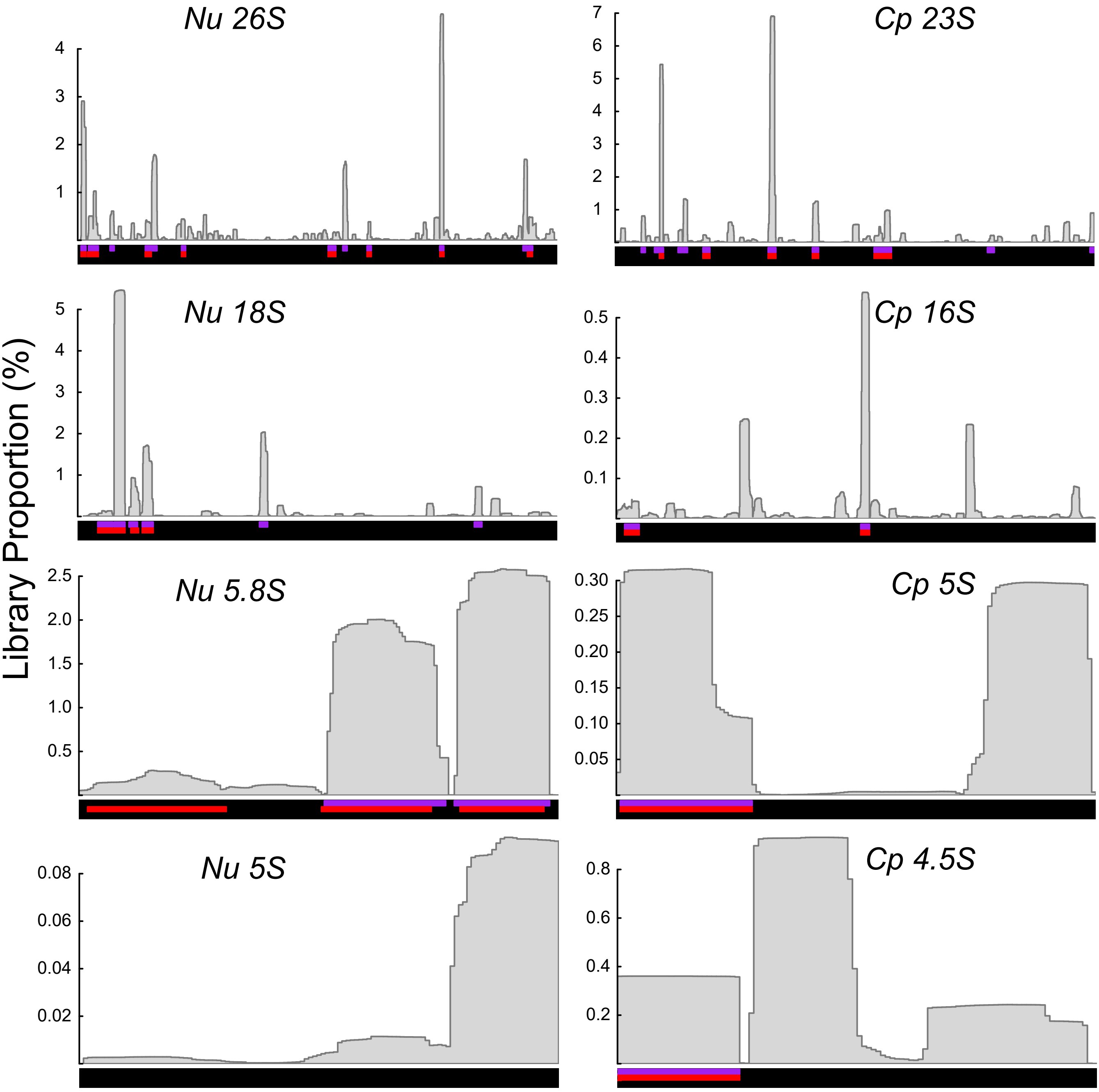

#### Cytosolic RPFs: Start codon

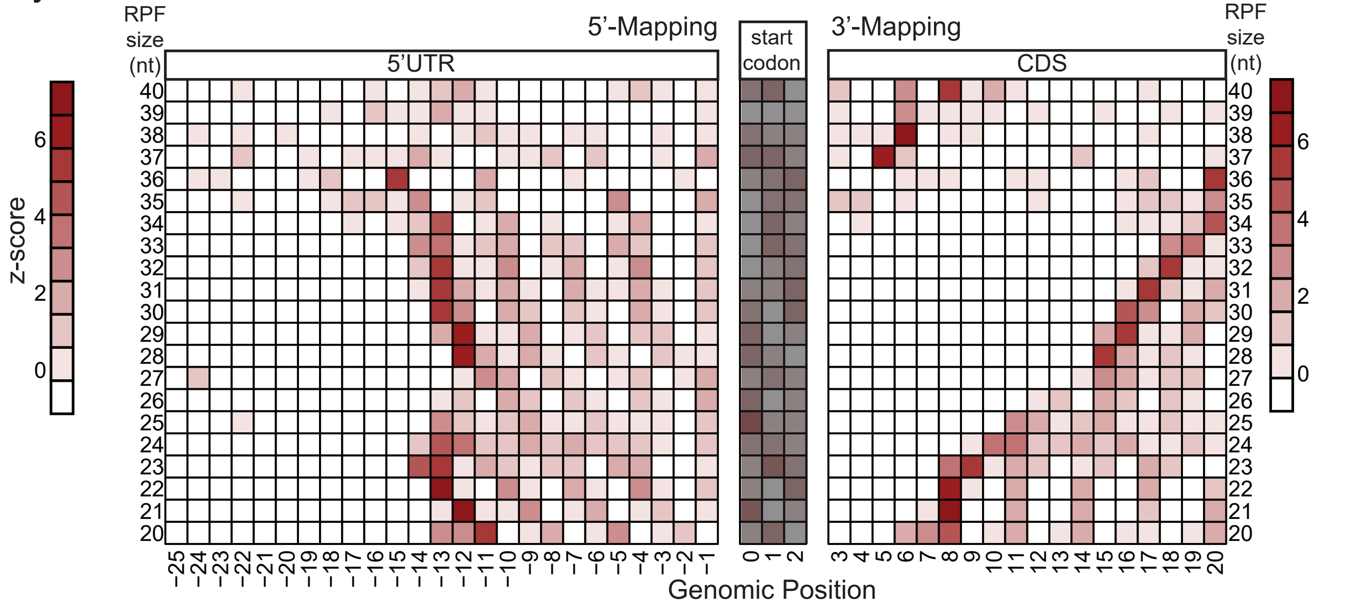


Figure S5
